# Essential PBP1-associated proteins of *Trypanosoma brucei*

**DOI:** 10.1101/2020.03.02.973057

**Authors:** L. Nascimento, M. Terrao, KK. Marucha, B. Liu, F. Egler, C. Clayton

**Affiliations:** Heidelberg University Centre for Molecular Biology (ZMBH), Im Neuenheimer Feld 282, D69120 Heidelberg, Germany

**Keywords:** Trypanosoma, mRNA, MKT1, PBP1, Ataxin-2, poly(A) binding protein

## Abstract

Control of gene expression in kinetoplastids depends heavily on RNA-binding proteins that influence mRNA decay and translation. We previously showed that MKT1 interacts with PBP1, which in turn recruits LSM12 and poly(A) binding protein. MKT1 is recruited to mRNA by sequence-specific RNA-binding proteins, resulting in stabilisation of mRNA. We here show that PBP1, LSM12 and an additional 117-residue protein, XAC1 (Tb927.7.2780), are present in complexes that contain either MKT1 or MKT1L (Tb927.10.1490). All five proteins are present predominantly in the complexes, and there was evidence for a minor subset of complexes that contained both MKT1 and MKT1L. MKT1 appeared to be associated with many mRNAs, with the exception of those encoding ribosomal proteins. XAC1-containing complexes reproducibly contained RNA-binding proteins that were previously found associated with MKT1. In addition, however, XAC1- or MKT1-containing complexes specifically recruit one of the six translation initiation complexes, EIF4E6-EIF4G5; and yeast 2-hybrid assay results indicated that MKT1 interacts with EIF4G5. The C-terminus of MKT1L resembles MKT1: it contains MKT1 domains and a PIN domain that is probably not active as an endonuclease. MKT1L, however, also has an N-terminal extension with regions of low-complexity. Although MKT1L depletion inhibited cell proliferation, we found no evidence for specific interactions with RNA-binding proteins or mRNA. Deletion of the N-terminal extension, however, enabled MKT1L to interact with EIF4E6. We speculate that MKT1L may either enhance or inhibit the functions of MKT1-containing complexes.

## Introduction

Transcription initiation is a crucial point of gene expression regulation in most eukaryotes (1). However, in kinetoplastid protists such as trypanosomes and leishmanias, transcription by the RNA polymerase II is polycistronic (2, 3), and mRNA precursors are co-transcriptionally processed to mature mRNAs by *trans*-splicing and polyadenylation (4, 5). To overcome the limitations imposed by such a system, the most abundant mRNAs are often encoded by multi-copy genes, or are even, in the case of trypanosome major surface proteins, transcribed by RNA polymerase I (6). Translation and decay of mRNAs are also at least partially linked: optimal codon usage correlates positively with mRNA stability, enabling the correct steady state levels of mRNAs and proteins from constitutively-expressed genes to be maintained (7, 8). However, many genes also require more flexible controls, allowing expression to be modified during differentiation and in response to environmental stimuli. For example, the African trypanosome *Trypanosoma brucei* alternates between two hosts during its life cycle, living in the blood and tissue fluids of mammals and the digestive system of tsetse flies. Adaptation to various niches within the two hosts is accompanied by changes in levels and translation of several hundred mRNAs (9–14). Regulated control of mRNA processing, translation and mRNA decay relies mainly on RNA binding proteins, which often (although not always) bind to the 3’-untranslated regions (3’-UTRs) of mRNAs (15). miRNAs appear to be absent (16).

Many RNA-binding proteins have been reported to control specific aspects of trypanosome gene expression (15). However, the only ones for which the molecular mechanism is understood are the zinc finger proteins ZC3H11 and ZC3H20. ZC3H11 binds to the 3’-UTRs of mRNAs that are required for survival after heat shock (17), while ZC3H20 binds to some mRNAs that are expressed specifically in procyclic forms (18, 19).

Stabilisation of mRNAs by ZC3H11 and ZC3H20 involves recruitment of a protein complex that includes MKT1, PBP1, LSM12 and poly(A) binding proteins (PABPs), especially PABP2 (18, 20). The trypanosome MKT1, PBP1 and LSM12 proteins were named after their previously-characterised *Saccharomyces cerevisiae* homologues Mkt1, Pbp1 and Lsm12 (21). In both organisms, Pbp1/PBP1 recruits all of the other subunits as well as PABP. The metazoan Pbp1 homologue, Ataxin-2, also interacts with both poly(A) binding protein and Lsm12 (22, 23), but metazoans lack Mkt1. Pbp1 and Ataxin-2 are incorporated into RNA-protein stress granules (24), as is *T. brucei* Pbp1 (25); and polyglutamine tract expansions in Ataxin-2 are implicated in neurodegenerative disease (22, 23). A point mutation in Mkt1 in the standard *S. cerevisiae* strain S288C has been linked to numerous minor defects (26–36); but the *T. brucei* MKT1 protein is wild-type at this position.

To assess the effects of proteins on bound RNAs in *T. brucei*, we use a tethering assay (37). We use a cell line that constitutively expresses a reporter mRNA bearing 5 copies of the lambda boxB sequence in the 3’-UTR. The boxB sequence is bound with high affinity by a short peptide from the lambdaN protein. We inducibly express proteins of interest, fused N-terminally to the λN peptide, in order to “tether” the fusion protein to the reporter RNA. Expression of λN-PABP, λN-PBP1, λN-LSM12, λN-MKT1, or λN-ZC3H11 all caused increases in reporter expression (20). Attachment of PABPs to mRNA 3’-UTRs increases mRNA levels (38–40), and this probably explains the mRNA-stabilising effects. Pull-down followed by mass spectrometry, as well as yeast two-hybrid analyses, showed that *T. brucei* MKT1 interacts with numerous RNA-binding proteins, often (though not always) via the motif (Y/W/V/I)(R/T/Q)H(N/D)PY (25). Such RNA-binding proteins can therefore stabilize their target mRNAs, and/or promote their translation, by recruiting the MKT1-PBP1-LSM12-PABP complex (18, 25).

We adapted the tethering assay in order to conduct a genome-wide screen for proteins that could influence mRNA fate (41). One of the strongest activators of gene expression was the protein encoded by Tb927.7.2780 (41). In this paper we show that this protein, which we name eXpression ACtivator 1 (XAC1), is present in two alternative complexes, containing PBP1, LSM12 and either MKT1 or an MKT1-like protein, MKT1L (Tb927.10.1490). We also find that both MKT1 and MKT1L are present mostly in complexes and that MKT1, but not MKT1L, is associated with many mRNAs and with a specific eIF4F like complex.

## Results

### XAC1 is essential for trypanosome survival

*T. brucei* XAC1 is 117 residues long and has no annotated functional domains (Figure 1A). Syntenic homologues are present in all kinetoplastid parasite genomes examined, but not in the free-living *Bodo saltans* or evolutionarily more distant organisms. A 17-residue region is conserved in all sequences examined (Figure 1B and black box in Figure 1A), and the sequences from *Trypanosoma* species have two regions rich in proline, alanine, glutamine and serine further towards the N-terminus (marked red in Figure 1).

**Figure 1.**
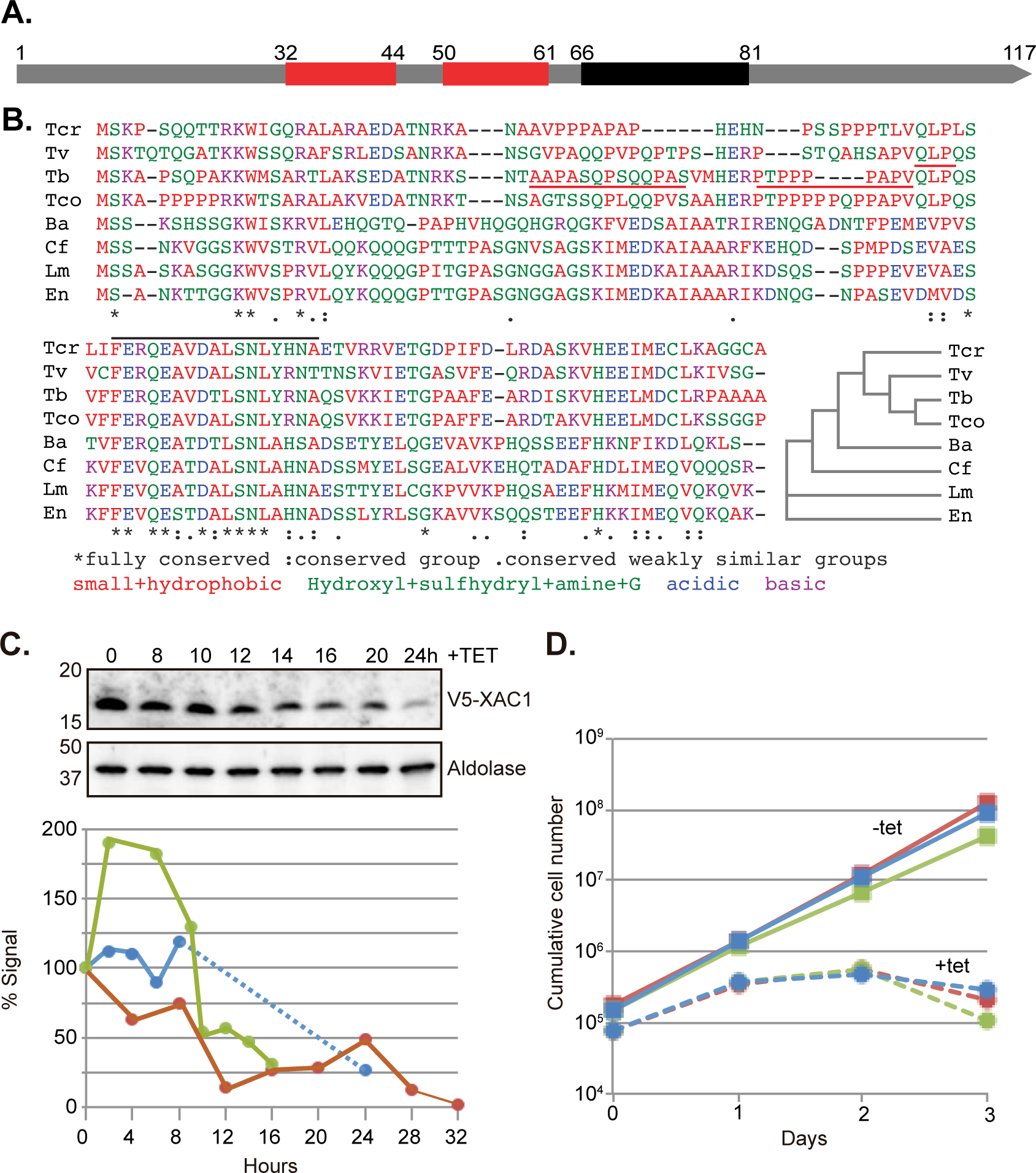
XAC1 is essential for cell survival. A) Schematic representation of *T. brucei* XAC1. The black rectangle indicates a region that is conserved in all parasitic kinetoplastids, and the red rectangles indicate regions of relatively low complexity in the *T. brucei* sequence. Amino acid positions are indicated. B) Multiple sequence alignment of XAC1 from various kinetoplastid species using Clustal Omega (EMBL-EBI). Tcr -*Trypanosoma cruzi* TcCLB.510525.40; Tv - *Trypanosoma vivax* TvY486_0043210; Tb - *T. brucei* Tb927.7.2780; Tco - *Trypanosoma congolense* TcIL3000_7_2100; Ba - *Blechomonas ayalai* Baya_018_0640; Cf - *Crithidia fasciculata* CFAC1_240015600; Lm - *Leishmania major* LmjF.22.0890; En - *Endotrypanum monterogeii* EMOLV88_220012800. The low complexity regions are indicated by red lines and the most conserved region with a black line. An evolutionary tree is on the right. C) XAC1 quantification upon RNAi induction in bloodstream forms. Cells with *in situ* V5-tagged XAC1 were made by integrating a PCR product. The V5 signal was quantified by Western blot over 16-24 hours of RNAi induction. Aldolase was used as a loading control. Top panel: western blot; bottom panel: blot quantification (by ImageJ) for 3 independent experiments, shown in different colors. D) Growth curve of V5-XAC1 expressing bloodstream-form trypanosomes upon RNAi induction. Cumulative cell numbers are given for three different clones after various days of incubation with or without tetracycline.

The results of a high-throughput RNAi screen suggested that XAC1 is essential for trypanosome growth (42). In order to be able to monitor XAC1 protein levels we integrated a sequence encoding a V5 tag immediately upstream of, and in frame with, an *XAC1* open reading frame. Stem-loop RNAi induction (Supplementary Table S1) resulted in a 75% reduction of V5-XAC1 protein within 16h (Figure 1C). At this point, protein synthesis was still normal (Supplementary Figure S1A), but no subsequent growth was seen (Figure 1D).

### Composition of XAC1-containing complexes

To investigate XAC1 interactions, we first made bloodstream-form trypanosomes in which one *XAC1* gene included C-terminal tag for tandem affinity purification (43) (XAC1-TAP), and the other *XAC1* gene had been deleted (Supplementary Table S1). Results for GFP-TAP served as negative controls; some of those results have been published previously (44, 45). Results from three tandem affinity purifications of XAC1-TAP are shown in Figure 2A and Supplementary Table S2. The most strongly enriched proteins were MKT1, LSM12, PBP1, Poly(A) binding protein 2 (PABP2), and a protein related to MKT1, which we named MKT1L (Tb927.10.1490) (Figure 2A). Digestion with RNase (not shown) did not affect this. We separately confirmed pull-down of YFP-tagged XAC1 by myc-tagged LSM12 (Supplementary Figure S2A).

**Figure 2.**
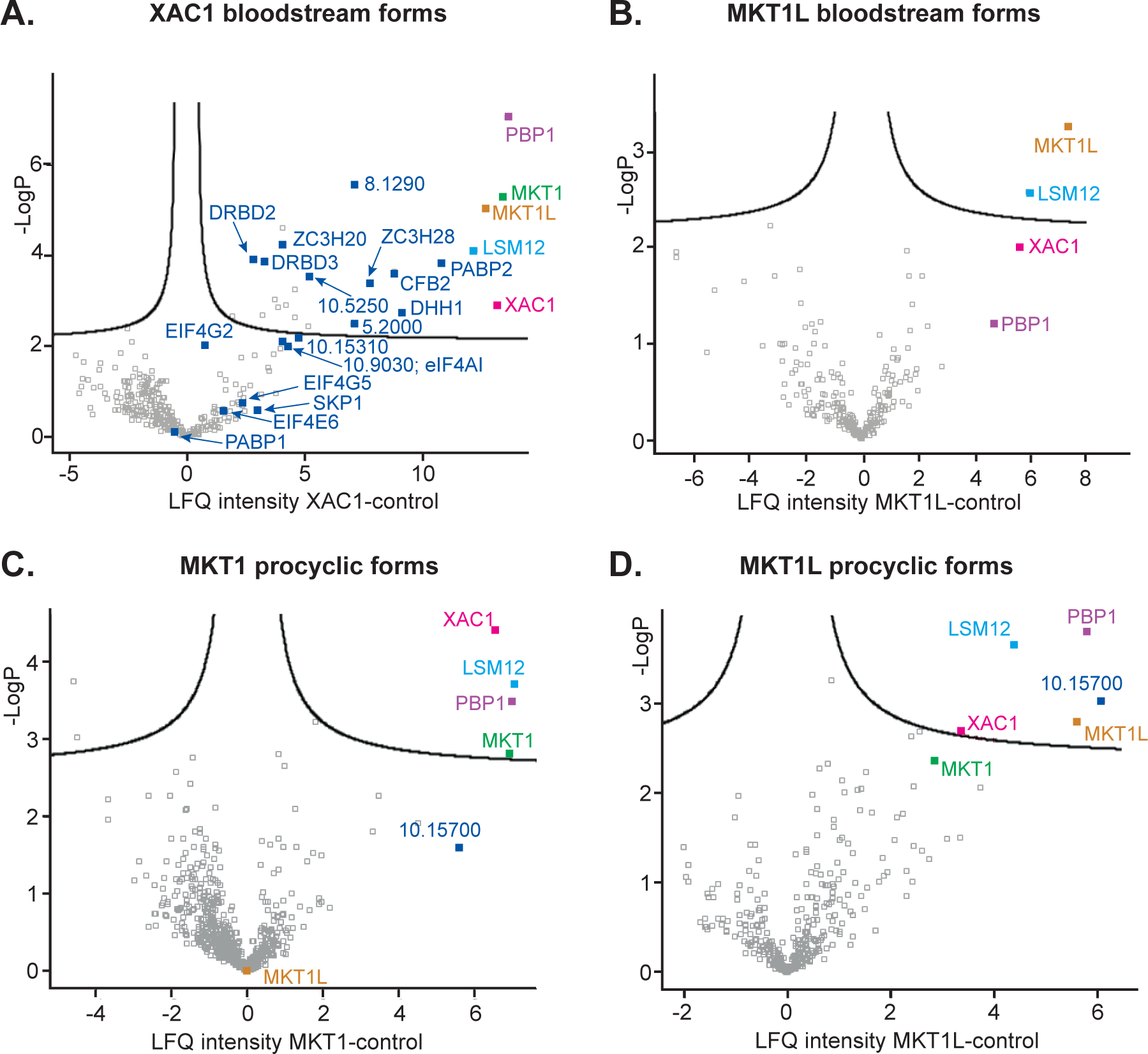
Proteins that co-purify with XAC1, MKT1 and MKT1L. (A) Volcano plot of bloodstream forms XAC1 interactors. The *XAC1* gene was tagged *in situ* with a tandem affinity purification tag (protein A - TEV cleavage site - calmodulin binding peptide) and the other gene was replaced by a selectable marker (*NPT*). After two cycles of purification the preparation was analysed by mass spectrometry. GFP-TAP served as the control. Detailed analysis using MaxQuant and PERSEUS was as described in the Methods section. The identities of selected spots are indicated; all significantly enriched proteins are listed in Table 1, and details are in Supplementary Table S2. The colour code is as in Figure 3. (B) Volcano plot of MKT1L interactors in bloodstream forms. MKT1L-TAP was inducibly expressed in bloodstream forms and purified as in (A). For details of the analysis, see Supplementary Table 2. (C) Volcano plot of procyclic form MKT1 interactors. Procyclic forms with the *MKT1* gene tagged *in situ* with a V5 tag were used. The V5-MKT1 protein was immunoprecipitated, then the preparation was analysed by mass spectrometry. V5-GFP (pHD3141) served as the control. For details of the analysis see Supplementary Table 2. (D) Volcano plot of procyclic form MKT1L interactors. Procyclic forms with the *MKT1L* gene tagged *in situ* with a V5 tag (pHD 2730) were used. the V5-MKT1L protein was immunoprecipitated, then the preparation was analysed by mass spectrometry. V5-GFP served as the control. For details of the analysis see Supplementary Table 2.

Previously, we had identified proteins associated with MKT1 in bloodstream forms after a single purification of C-terminally TAP-tagged MKT1, followed by non-quantitative mass spectrometry. We had also found interaction partners in a yeast 2-hybrid screen (25). In addition to PBP1, LSM12, MKT1L and MKT1, 24 proteins were significantly enriched with XAC1-TAP. Of these, ten had previously been identified as direct or indirect partners of MKT1 (25) (Table 1). Most of the additional XAC1-associated proteins were found cross- linked to poly(A)+ RNA (the mRNP proteome) (46), and they included seven with canonical RNA-binding domains. Beyond the main complex, nine of the associated proteins were also shown to activate expression when tethered (46, 47) (Table 1), consistent with an ability to recruit the MKT1-PBP1 complex.

**Table 1.** Proteins from bloodstream-form trypanosomes that co-purify with XAC1. Columns are as follows: 1:- Tb927 gene ID. 2:- protein name: UF-unknown function, 3:- subcellular location ((54) or other data): C-cytoplasm, N-nucleus, Flag-flagellum, UF-unknown function. 4:- presence in the mRNA proteome (46): yes - False Discovery Rate (FDR) <0.05, maybe - FDR >0.05, No - not detected, Yes* - ZC3H20 has RNA-binding domains and binds to mRNAs in procyclic forms (19), but has low abundance in bloodstream forms so was not in the mRNP proteome. 5:- effect of tethering to a reporter: A- activates as a full-length protein (46), (A) activated in a high-throughput screen of protein fragments (47), *shown or confirmed in this work; (a) has been shown by other means to stabilise its target mRNAs (55); R-represses as full-length protein, 0-no effect detected. 6- % coverage in a single purification of MKT1-TAP (25). 7:- interaction with MKT1 in yeast-two hybrid assay (individual or screen) - number of different interacting fragments detected (25). 8:- log10 P-value for co-purification with XAC1-TAP. 9:- Enrichment with XAC1-TAP. Values for (8) and (9) were obtained using PERSEUS.

| 1 | 2 | 3 | 4 | 5 | 6 | 7 | 8 | 9 |
| --- | --- | --- | --- | --- | --- | --- | --- | --- |
| GeneID | What | Location | mRNP | Tether | MKT1-TAP | MKT1 Y2H | -Log pvalue | Enrichment |
| 6.4770 | MKT1 | C | Yes | A | 69% | 0 | 5.3 | 13.4 |
| 10.14900 | MKT1L | C | Yes | A | 20% | 0 | 5.0 | 12.6 |
| 9.9060 | LSM12 | C | Yes | A | 60% | 0 | 4.1 | 12.2 |
| 7.2780 | **XAC1** | C | No | A* | 71% | 0 | 2.9 | 13.1 |
| 8.4540 | PBP1 | C | Yes | A | 45% | 8 | 7.1 | 13.6 |
| 9.10770 | PABP2 | C | Yes | A | 0% | 0 | 3.9 | 10.8 |
| 1.4650 | CFB2 | C | Yes | A | 40% | 9 | 3.7 | 8.8 |
| 7.2660 | ZC3H20 | C | Yes* | (A) | 0% | 11 | 4.3 | 4.0 |
| 9.9450 | ZC3H28 | C | Yes | (A) | 0% | 0 | 3.4 | 7.8 |
| 10.5250 | ZC3H32 | C | Yes | R | 5% | 16 | 3.5 | 5.2 |
| 10.11760 | PUF6 | C | Yes | A | 0% | 6 | 3.0 | 3.7 |
| 1.2600 | PUF9 | C | Yes | (a) | 0% | 7 | 2.7 | 3.2 |
| 9.13990 | DRBD2 | C | Yes | R | 0% | 0 | 3.9 | 2.9 |
| 9.8740 | DRBD3/PTB1 | C | Yes | (A) | 0% | 0 | 3.9 | 3.3 |
| 10.3990 | DHH1 | C | Yes | 0 | 10% | 0 | 2.7 | 9.1 |
| 10.14550 | DED1-1 | C | Yes | 0 | 0% | 0 | 2.6 | 4.6 |
| 3.3180 | NUP98 | N | No | 0 | 0% | 0 | 4.6 | 4.1 |
| 2.4230 | NUP1 | N | No | 0 | 0% | 0 | 2.9 | 4.2 |
| 10.8940 | KHARON1 | Flag | Yes | 0 | 0% | 0 | 2.7 | 1.9 |
| 8.1290 | UF | C | Maybe | 0 | 0% | 8 | 5.6 | 7.1 |
| 5.2620 | UF | C | Yes | (A) | 0% | 0 | 3.3 | 4.6 |
| 11.3440 | UF | C | Yes | A | 9% | 2 | 2.4 | 5.2 |
| 5.200 | UF | C | Yes | 0 | 28% | 0 | 2.5 | 7.1 |
| 8.3850 | UF | C | Yes | (A) | 0% | 3 | 2.9 | 3.9 |

### Structure and protein interactions of MKT1L

MKT1L has annotated “Mkt1-like” PIN, N- and C-terminal domains (E-values 1e-22, 1e-15 and 2 e-14 respectively) (Figure 3A), but there is only 20% identity with MKT1 over the Mkt1 domain-containing region (Supplementary Figure S3). MKT1L is distinguished from MKT1 by an extended N-terminus that includes 35- residues consisting almost exclusively of glutamine and histidine, which are predicted to be disordered by mobibd_LITE. Similar low-complexity regions are found in the *Trypanosoma cruzi* and *Trypanosoma congolense* homologues, and the region is expanded to 130 residues in *Trypanosoma vivax* (Supplementary Figure S3). The low-complexity region is followed by a sequence that, in 2018, was classified as having a weak resemblance to the Med15 domain (Pfam 09606) but this was no longer recognised as significant in the 2020 version of Prosite. The function of Med15 domains is not known.

**Figure 3.**
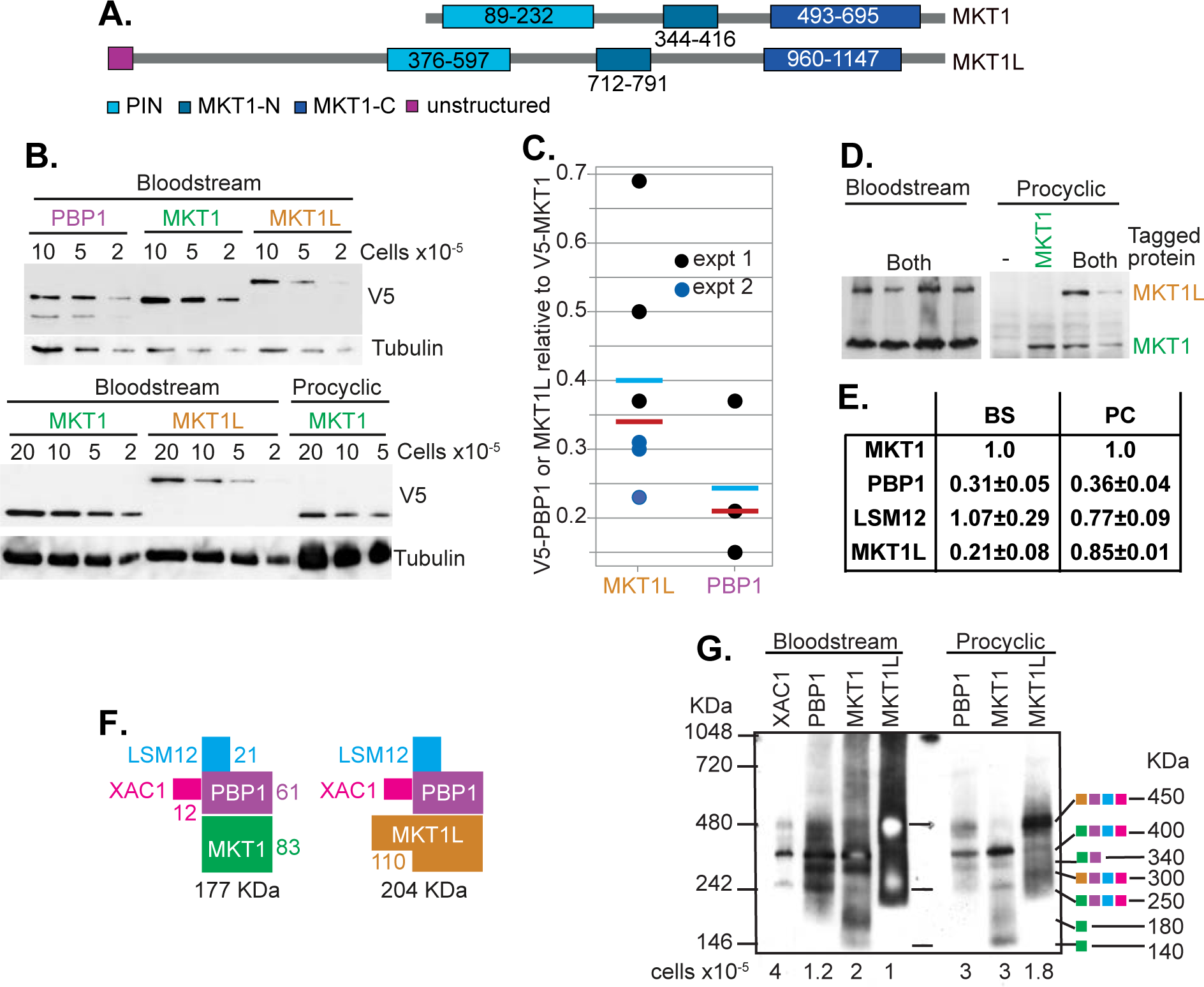
Quantitation and complex composition. (A) Comparison of MKT1 and MKT1L, to scale. The different domains and their positions in the sequence are labelled. (B-D) Trypanosomes with *in situ* tagged MKT1 (pHD1973), MKT1L (pHD 3071) or PBP1 (pHD2166) were pelleted and boiled in SDS sample buffer. The abundances were then measured by Western blotting, with overnight transfer to allow detection of high-molecular weight proteins. Signal intensities were assessed using ImageQuant and the resulting graph is shown below. (“Signal” is arbitrary grey-scale units). (B) Western blot data for panel (B). Experiment 2 is above and experiment 1 below. BS-bloodstream forms; PC-procyclic forms. The numbers of cells used per lane, multiplied by 10^-5^, are shown. (C) Amounts of V5-PBP1 and V5-MKT1L relative to V5-MKT1 in bloodstream forms. Individual results for different dilutions are shown. The cyan bars are arithmetic mean and the magenta bars are the median. (D) Results for cells in which both MKT1 and MKT1L were tagged. (E) Levels of PBP1, LSM12 and MKT1L relative to MKT1 in total proteomes of Bloodstream (BS) and Procyclic (PC) forms. The Label Free Quantification (LFQ) values from four replicate published proteomes for each stage (48) were used to calculate means and standard deviations of the ratios. XAC1 was not reproducibly detected in bloodstream forms, perhaps because of its small size. These values are not reliable by themselves because some peptides are more readily detected than others, but the increased relative amount of MKT1L in procyclic forms is clear. (F) Diagrams of two possible complexes. (G) Blue native gel electrophoresis of extracts from bloodstream and procyclic forms expressing V5-tagged XAC1, PBP1, MKT1 or MKT1L was followed by Western blotting and detection of the V5 tag. The numbers of loaded cell equivalents are indicated below the blot. Proteins detected in individual bands, with their measured complex molecular weights, are indicated on the right using the colour code in panel F.

In our previous studies of MKT1 interactions, we had used a C-terminal TAP tag. To confirm the interaction of MKT1L with XAC1, we therefore similarly inducibly expressed MKT1L-TAP and purified associated proteins. Rather to our surprise, the only proteins that co-purified were components of the known MKT1 complex: LSM12, and XAC1, with PBP1 enriched but below the significance cut-off (Figure 2B, Supplementary Table S2). No RNA-binding proteins were associated, even under more relaxed parameters. These results, combined with previous data, suggest that the many additional proteins that were found to be associated with XAC1 were probably attached via MKT1, not MKT1L.

We next examined the proteins associated with MKT1 and MKT1L in procyclic forms. Since effects of the large C-terminal tag, or of inducible (over)-expression could not be ruled out, we decided to use cell lines in which one of the relevant genes contained a sequence encoding a V5 tag at the 5’-end of the open reading frame (*in situ* V5 tag), using V5-GFP expressed constitutively from the tubulin locus served as a negative control. This allowed for only a single round of purification, but this can be and advantage when quantitative mass spectrometry is used, since “background” sticky proteins can be measured in the negative control. Otherwise, they re simulated at random, and this can even result in proteins that are present in every test preparation, and completely absent in the control, being classified as non-significant (see “exclusive” in Supplementary table S2). The results were not as we had hoped. MKT1 from procyclic forms indeed co-purified XAC1, LSM12 and PBP1 (Figure 2C), but there was no significant enrichment of any RNA-binding proteins (Supplementary Table S2), perhaps because RNA-bidning proteins will all be present in sub-stoichiometric amounts and the background was now too high. MKT1L was robustly associated with XAC1, LSM12 and PBP1, with some MKT1 also detected (Figure 2D) (Supplementary Table S2). MKT1L also pulled down a 350 kDa repetitive protein (Tb927.10.15700) implicated in intra-flagellar transport, but the significant enrichment may be an artefact caused by the numerous repeats. These results at least, however, confirmed that the complexes were present in procyclic forms.

### MKT1, XAC1 and MKT1L are present only as complexes

Our mass spectrometry results clearly showed the existence of two complexes: MKT1-XAC1-PBP1-LSM12, and MKT1L-XAC1-PBP1-LSM12. LSM12 and MKT1 are known to interact with PBP1 (25). Quantitative mass spectrometry results are affected by the degree to which the peptides are detected, but if taken as approximate indicators, they suggested that each complex contained roughly equimolar amounts of XAC1, LSM12, PBP1 and either MKT1 or MKT1L. From the mass spectrometry results, it was clear that MKT1 and MKT1L were mostly present in different complexes. However there was some evidence for complexes containing both proteins. MKT1L had been detected after purification of MKT1 from bloodstream forms (25), but was not enriched after pull-down of V5-MKT1 from procyclic forms; MKT1 was not detected after purification of MKT1L-TAP from bloodstream forms, but was present after pull-down of V5-MKT1L from procyclic forms (Figure 2). To investigate complexes in more detail we used cell lines with *in situ* V5-tagged genes. By tagging at the N-terminus, changes in regulatory 3’-untranslated region sequences are avoided; however the change in the N-terminus of the protein might affect protein stability.

We first analysed protein abundance by denaturing polyacrylamide gels and quantitative Western blotting of the *in situ* tagged versions of the desired proteins. For bloodstream forms, quantitation of the signals suggested that both MKT1L and PBP1 are 2-4-fold less abundant than MKT1 (Figure 3B, C, D). Procyclic forms had less MKT1, with signals similar to those from MKT1L (Figure 3C, D). These results are consistent with published quantitative mass spectrometry data (Figure 3E) (48). The total proteome of procyclic forms has roughly equal amounts of MKT1 and MKT1L, while in bloodstream forms MKT1 is about 3 times more abundant than the MKT1L. These results are all consistent with competition between MKT1 and MKT1L for PBP1.

A complex containing MKT1, PBP1, XAC1 and MKT1 has a predicted mass of 177KDa, while a similar complex with MKT1L would be 204 KDa (Figure 3F). Results from blue native gel electrophoresis are shown in Figure 3G. Strangely, the signal from MKT1L for bloodstream-form extracts was much stronger than for the other proteins investigated. Perhaps MKT1L complexes were easier to solubilise, and some MKT complexes are in complexes that are too large to enter the gel - such as polysomes or aggregates? We do not know whether the migration of the complexes in the gel reflects their molecular weights with any degree of accuracy. If so, the interpretations in Figure 3G are possible. The predominant band for XAC1, PBP1 and MKT1 migrated at 400 KDa; this might be a MKT1-XAC1-PBP1-LSM12 dimer. Similarly, the equivalent MKT1L complex, at 450 KDa, would to be predominantly dimeric. Equivalent monomeric complexes may be represented by the bands that migrate at 250 and 300 KDa respectively.

It was notable that there was no MKT1L, PBP1 or XAC1 signal at apparent molecular weights below 250 KDa, suggesting that these three proteins are stable only within complexes. The presence of a PBP1 degradation product in the denaturing gels (Figure 3C) is consistent with this. MKT1 showed native gel signals at 140KDa and 180KDa (Figure 3G); these might be MKT1 multimers, or might represent MKT1 association with other proteins that were not examined, such as PABP2 or the multifunctional RNA helicase DHH1, which was also relatively abundant in the XAC1 purification. There was a very faint MKT1 signal at about 450 KDa (clearer in the results for procyclic forms). This might come from complexes containing both MKT1-XAC1-PBP1-LSM12 and MKTL-XAC1-PBP1-LSM12. Such heterodimers might explain our discovery of MKT1L after purification and mass spectrometry of MKT1-TAP from bloodstream forms (25), and MKT1 after purification of V5-MKT1L from procyclic forms (Figure 2D).

We concluded that MKT1 and MKT1L form separate complexes with PBP1 and XAC1, but that complexes containing MKT1 and MKT1L may also rarely associate with each other.

### Interactions of MKT1 and MKT1L with EIF4G5-EIF4E6 complexes

To investigate interactions within MKT1 and MKT1L complexes we used yeast two-hybrid assays. No interaction of XAC1 with MKT1 was detected. PBP1 (25), XAC1 and MKT1L fused with the DNA-binding domain (bait) activated by themselves, so mutual interactions of these proteins could not be tested (Supplementary Figure 5A, B). Deletion of the repetitive N-terminus of MKT1L, however, eliminated the self-activation; and this protein interacted with PBP1 (MKT1LΔN, Figure 4A, B, and Supplementary Figure S4). We therefore concluded that MKT1L, like MKT1, binds to PBP1. The means by which XAC1 is associated with the complex is unresolved.

**Figure 4.**
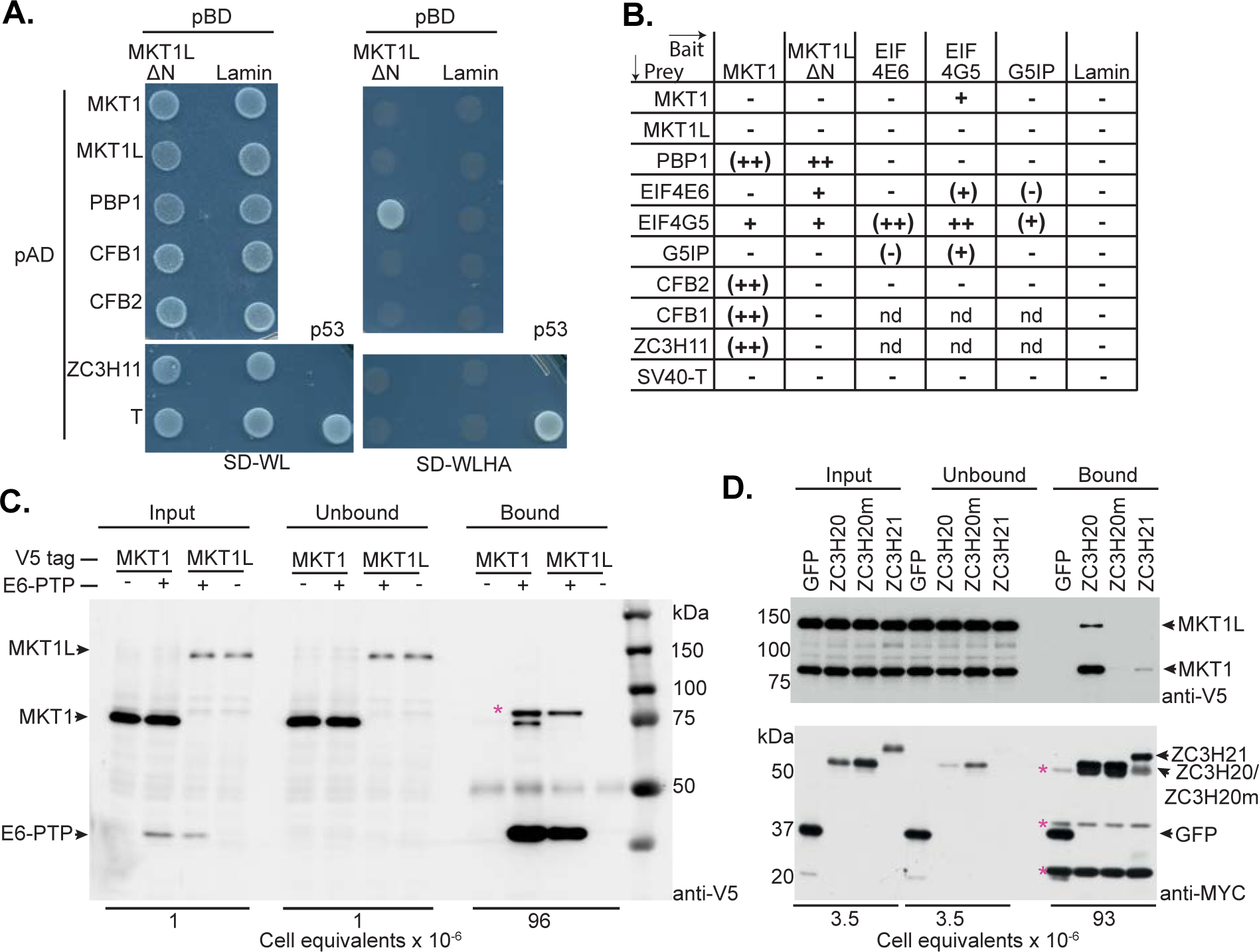
Interactions of XAC1, PBP1 and MKT1L in a yeast two-hybrid assay. (A) Interactions of MKT1LΔN with known MKT1 interactors. The *Saccharomyces cerevisiae* AH109 strain was transformed with the indicated plasmid combinations of bait and prey. Transformed yeast cells were grown on selective media lacking Ade, His, Leu, Trp (Quadruple drop-out, QDO) to test protein interaction. The positive control on the bottom right is the interaction between SV40 T antigen (AD) and p53 (BD). (B) Conclusions from several replicates (see Supplementary Figures S4 and S5). Some of the proteins, including PBP1 (20), XAC1, CFB2 and MKT1L, are “self-activators” when fused with the DNA-binding domain, giving growth on the QDO media and variable beta-galactosidase staining independent of the co-expressed AD protein (e.g. MKT1L-BD interacting with the negative control, Lamin-AD). In parentheses, previously published interactions. (C) MKT1 associates with EIF4E6. The bloodstream-form cells used expressed *in situ* V5-tagged MKT1 or MKT1L, and *in situ* PTP-tagged EIF4E6. EIF4E6-PTP was pulled down on IgG magnetic beads and released by boiling. The anti-V5 antibody binds to the PTP tag. The asterisk indicates an additional band that is reproducibly seen in the EIF4E6-PTP line; we do not know what it is. (D) Interactions of ZC3H20 and ZC3H21 with MKT1 and MKT1L. The procyclic trypanosomes used expressed *in situ*-tagged V5-MKT1 and V5-MKT1L, and inducible myc-tagged GFP, ZC3H20, ZC3H20 with a HHYNPY -> HYNAA mutation, and ZC3H21. After immunoprecipitation with anti-myc beads the bound proteins were released by boiling and analysed by Western blotting. Asterisks indicate IgG chains.

Intriguingly, the translation initiation complex EIF4E6-EIF4G5 (49) was present in the XAC1 affinity purification (Figure 2A, Supplementary Table S2). The enrichment was below the significance cut-off, and the intensities were about 100-fold lower than for MKT1 complex components. However, neither EIF4E6 nor EIF4G5 was detected in the GFP-TAP control and the known abundant initiation complexes EIF4E3-G4 and EIF4E4-G3 (50) were undetectable in the XAC1 preparations. A pull-down of *in situ* PTP-tagged EIF4E6 indeed revealed V5-MKT1, but no association with V5-MKT1L was seen, even after prolonged exposure of the blot (Figure 4C and Supplementary Figure S2B). In two-hybrid assays, as expected, there was a strong interaction between EIF4E6 and EIF4G5, and between EIF4G5 with G5IP; there was also a novel interaction of EIF4G5 with itself. But notably, in addition, EIF4G5 interacted with MKT1 (Figure 4B and Supplementary Figure S4 and S5).

Interestingly, MKT1L did not interact with EIF4G5 neither EIF4E6, while the mutant MKT1LΔN did (Figure 4B and Supplementary Figure S4). Overall, these results suggest that EIF4E6-EIF4G5 complexes may be recruited to some mRNAs by MKT1.

### Do MKT1 and MKT1L interact with RNA-binding proteins in procyclic forms?

As noted above, associations of MKT1 with RNA-binding proteins were not detected in procyclic form extracts. In the bloodstream-form XAC1-TAP purifications, normalised peak intensities of significantly enriched RNA-binding proteins were 40-1000 times lower than that of MKT1, so perhaps even if they were somewhat enriched after a single purification step they would not be detected above the background. To check this, we decided to look at the interaction of MKT1 with ZC3H20 and ZC3H21. These two RNA-binding proteins have MKT1 interaction motifs and are more abundant in procyclic forms than in bloodstream forms (48). Moreover, the ability of ZC3H20 to activate when tethered in procyclic forms depends on the MKT1 interaction motif (18). ZC3H20 and ZC3H21 were detected in only one of the four V5-MKT1 preparations (Supplementary Table S2) although ZC3H20 was reproducibly detected in the bloodstream-form XAC1-TAP purification (Supplementary Table S2, Figure 2A). To test for interactions we used procyclic trypanosomes in which both MKT1 and MKT1L had *in situ* N-terminal V5 tags (Figure 3D). In these cells, we inducibly expressed GFP, ZC3H20, or ZC3H21 with C-terminal myc tags. Myc-tagged ZC3H20 pulled down V5-MKT1 and a little V5-MKT1L, while GFP, or a mutant version of ZC3H20 lacking the MKT1 interaction motif, did not (Figure 4D). Only V5-MKT1 was detected weakly in the ZC3H21-myc pull-down. These results suggest that ZC3H20 and perhaps ZC3H21 associate preferentially with MKT1. As expected, only a very small proportion of the MKT1 - about 1/30 - co-precipitated with ZC3H20.

So far we had very little evidence that MKT1L could interact with RNA-binding proteins. We therefore used the two-hybrid approach to test this directly for three proteins that are known to interact with MKT1 via C-terminal HNPY-like motifs: CFB1, CFB2, and ZC3H11. The N-terminally deleted version of MKT1L, which contains all MKT motifs, was indeed unable to interact with any of these proteins (Figure 4A, B). The yeast activation and DNA-binding domains were fused to the N-termini of the tested proteins, so the C-terminus of MKT1L was intact in these experiments. Although we cannot rule out an effect of the N-terminal deletion, this result further supports the conclusion that MKT1L does not directly associate with RNA-binding proteins that interact with MKT1.

### MKT1L is essential in bloodstream forms and procyclic forms

Results of an RNAi screen (51) suggested that MKT1L is an essential protein. Specific RNAi targeting MKT1L indeed resulted in strong growth inhibition in both bloodstream forms (Figure 5A, B) and procyclic forms (Figure 5C, D) within 1-2 days. The degree of growth inhibition in procyclic forms correlated with the extent of protein down-regulation (Figure 5C, D) and in one procyclic-form cell line, growth resumed after the RNAi effect was lost (Figure 5C,D). Protein synthesis was uniformly decreased 24h after MKT1L RNAi induction in bloodstream forms (Supplementary Figure S1B). This also happens after depletion of MKT1 (25) but it is not possible to tell whether the effect is direct or indirect. By Western blotting, the variant surface glycoprotein expressed by the trypanosomes without RNAi induction migrated as two bands (Figure 5B), and the slower-migrating band was reduced in abundance after MKT1L depletion. We do not know what this band is - it seems unlikely to represent recently synthesised protein with less glycosylation, since that is seen only by pulse-labelling (52).

**Figure 5.**
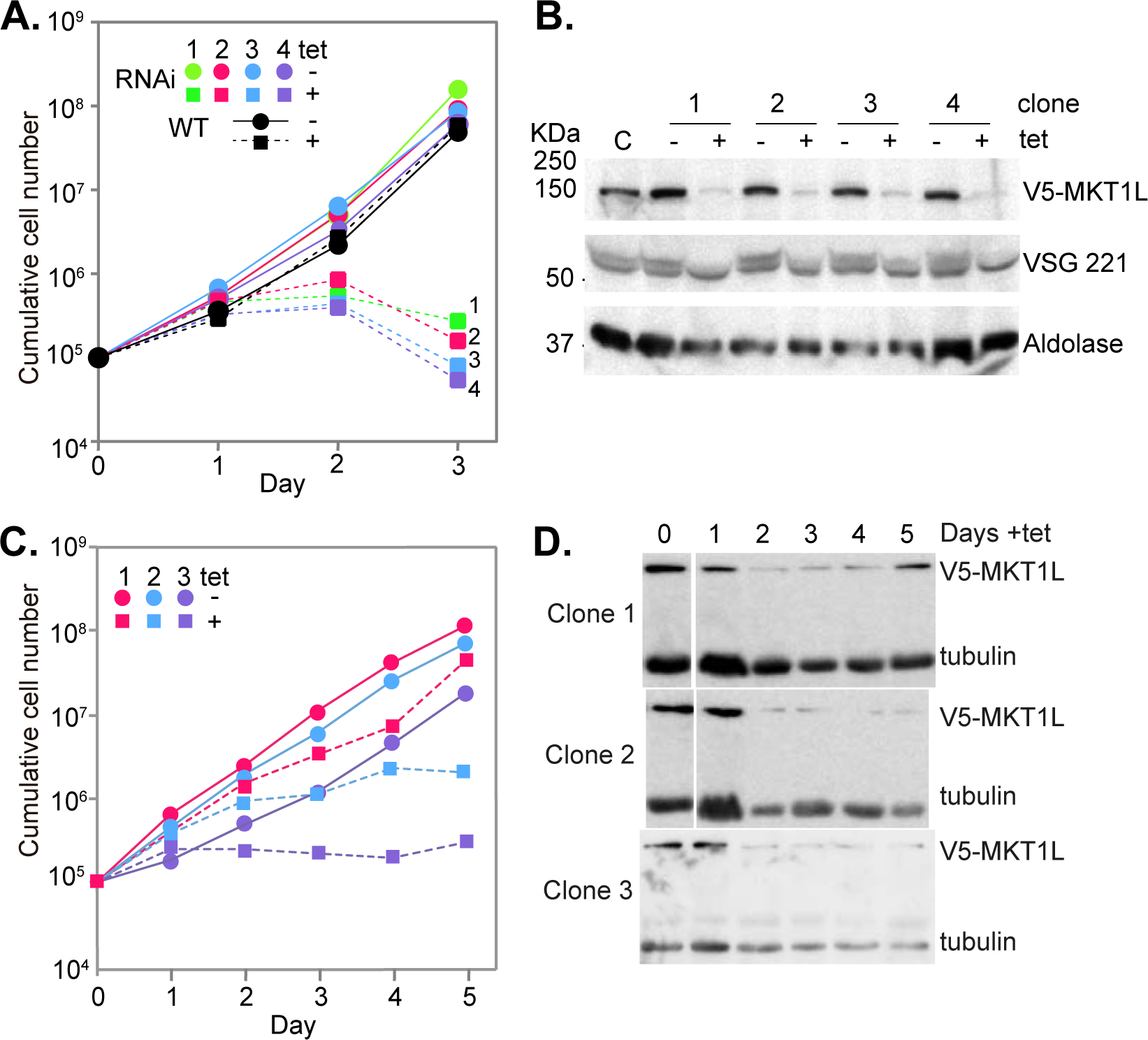
MKT1L is essential for survival. A) Cumulative growth of V5-MKT1L-expressing bloodstream forms with and without RNAi induction. The cultures were maintained in the presence (+tet) or absence (-tet) of tetracycline during 3 days. Control cells are cells expressing only the *Tet* repressor in the presence or absence of tetracycline. Cumulative cell numbers for four clones are shown. B) *MKT1L* RNAi induction in the four clones shown in (A). RNAi was induced for 24h in a cell line expressing *in situ* V5-MKT1L. The loading controls were VSG 221 and aldolase. C) Cumulative growth of V5-MKT1L-expressing procyclic forms with and without RNAi induction. Cultures were maintained for 5 days. Control cells are clones without addition of tetracycline. Cumulative cell numbers for three clones are shown. D) *MKT1L* RNAi induction in the three procyclic clones shown in (C). Tubulin was used as loading control.

We had previously found that MKT1 depletion was rapidly lethal in bloodstream forms, but had no effect in procyclic forms (25). We therefore attempted to delete both MKT1 genes in procyclic forms, using constructs bearing two different selectable markers (Supplementary Figure S6A). Although doubly-resistant trypanosomes were obtained, the MKT1 open reading frame persisted in all clones (Supplementary Figure S6B), suggesting that duplication of the locus was necessary to allow cell survival.

We also tested the functionality of the MKT1-TAP protein. In bloodstream forms expressing MKT1-TAP, deletion of one endogenous *MKT1L* gene was straightforward, but we were able to obtain only one cell line that lacked both wild-type copies of *MKT1L* (Supplementary Figure S6C). Those cells grew very slowly in the presence of tetracycline to induce MKT1L-TAP expression (doubling time of more than 24h), suggesting either that MKT1L-TAP is only partially functional, or that the level expressed was insufficient. Reduction of the inducer level to 2ng/ml or less, however, resulted in cell death, showing that MKT1L-TAP could at least partially substitute for MKT1L function.

### Locations of XAC1 and MKT1L

To check the subcellular locations of XAC1 and MKT1L we used cell lines expressing tagged versions. Both V5- and YFP-tagged XAC1 were in the cytoplasm (Figure 6A, B). In the cell line expressing *in situ* YFP-tagged XAC1, the second allele of *XAC1* was replaced by the *NPT* gene, demonstrating the functionality of the YFP-tagged version.

**Figure 6.**
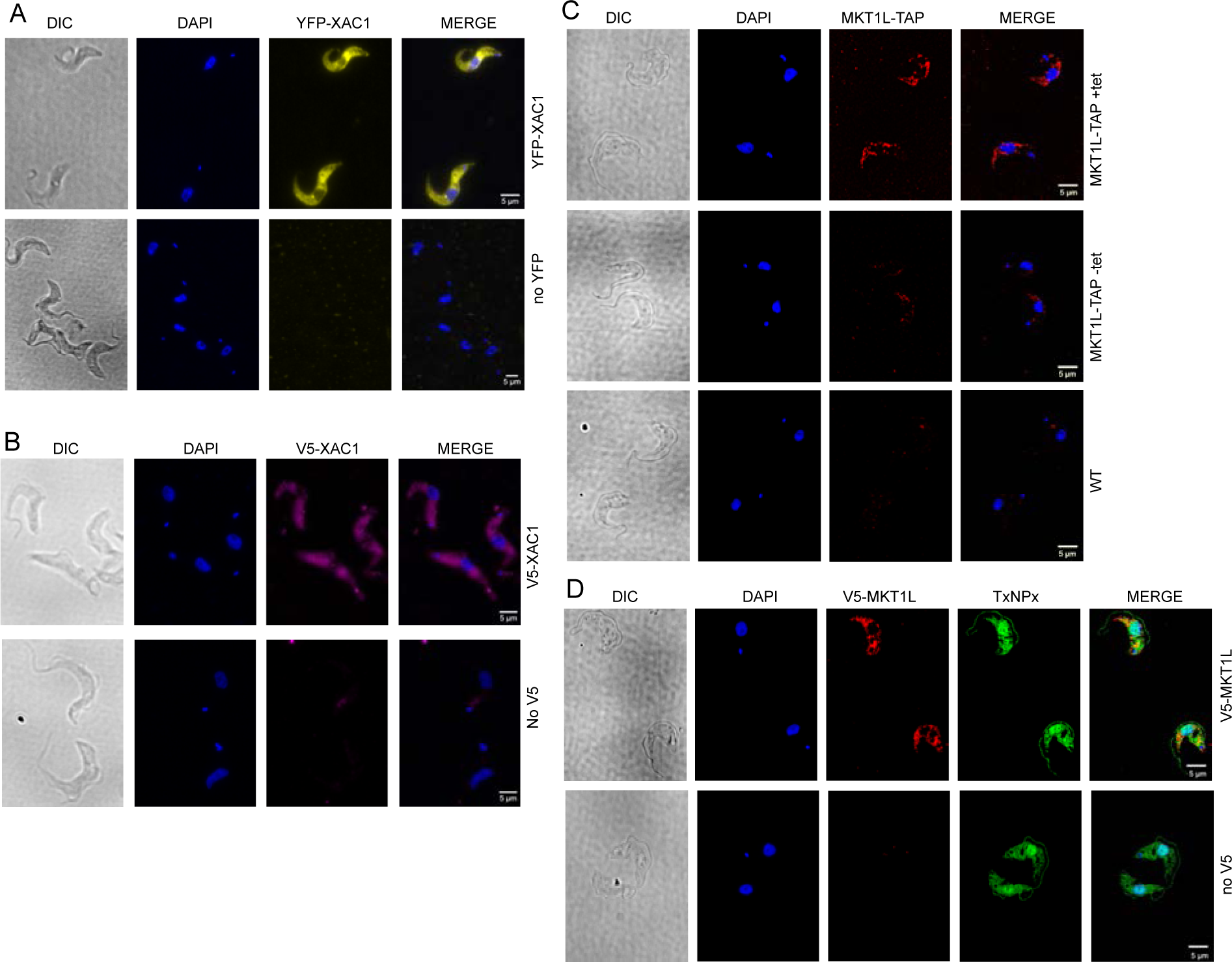
Tagged XAC1 and MKT1L are in the cytoplasm. In all preparations, the DNA (nuclei and kinetoplasts) was detected with diaminopimelic acid (DAPI). Differential interference contrast (DIC) images are on the left. All the images were acquired using an Olympus IX81 inverted Microscope. A) *In situ* tagged YFP-XAC1 detected by fluorescence microscopy using the YFP filter. The second copy of the protein in these transfectants was replaced by the *NPT* gene, demonstrating functionality of YFP-XAC1. B) *In situ* tagged V5-XAC1 detected by anti-V5 antibody. C) MKT1L-TAP detected by anti-PAP antibody. Induction was for 24 hours. D) *In situ* V5-MKT1L detected by anti-V5 antibody, with anti-tryparedoxin peroxidase (TxNPx) as a control.

MKT1L was previously found in *Leishmania tarentolae* splicing complexes that were purified using tagged SmD3 and U1A (53). Moreover, inducibly-expressed PTP-tagged MKT1L was found in speckles around the nuclear periphery of procyclic forms (53). These properties would not be consistent with a role of MKT1L in cytosolic mRNA control. We, however, detected no association of tagged *T. brucei* MKT1L with spliceosomal components by mass spectrometry. Moreover, both inducibly expressed MKT1L with a C-terminal TAP tag (Figure 6C) and MKT1L tagged *in situ* with an N-terminal V5 tag (Figure 6D) were in the cytoplasm and almost completely excluded from the nucleus of bloodstream forms. Resuts from the Tryp-tag project indicated that *in situ* C-terminally GFP-tagged MKT1L is spread throughout the cytoplasm of procyclic forms, with only a few faint nuclear spots (54) (http://tryptag.org/?id=Tb927.10.1490); transfection o obtain N-terminally GFP-tagged MKT1L were unsuccessful. Overall, the results suggest that most MKT1L is cytoplasmic.

### Effects of XAC1 and MKT1L in the tethering assay

We had previously shown that tethering of XAC1 increases expression of a *CAT-boxB* reporter, with an approximately 7-fold increase in CAT activity (41). In repeat assays we measured a 3 - 3.5 -fold increase in activity and a slightly lower increase in reporter mRNA (Figure 7A, Supplementary Figure S7). The results of a deletion analysis suggested that the entire protein was required for the full increase in CAT activity and mRNA, although the N-terminal residues might give a slight enhancement of translation (Figure 7A). Since MKT1L interacts with PBP1 we expected that it too would increase expression when tethered. Indeed, MKT1L tethering increased both CAT activity and mRNA (Figure 7B).

**Figure 7.**
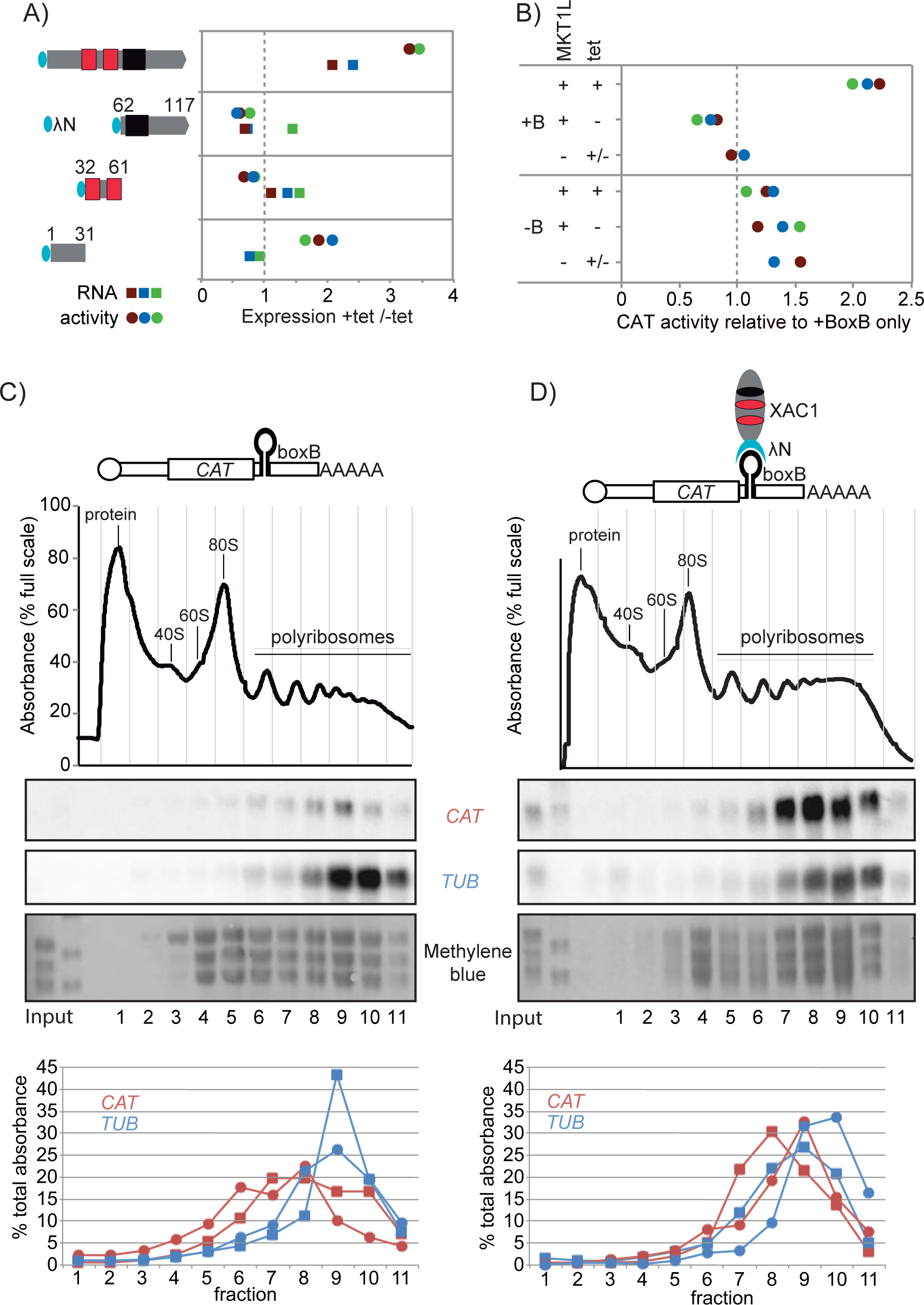
Tethered XAC1 and MKT1L increase expression from a reporter mRNA. A) Tethering assay using the XAC1 full length protein and fragments. XAC1 domains are as in Figure 1A. Levels of CAT activity and *CAT* mRNA in cell lines with inducible fusion protein expression were measured in the presence and absence of tetracycline. The ratios are shown in different colours for three independent cell lines. Expression of the tagged proteins is shown in Supplementary Figure S7A and B. B) Tethering assay with MKT1L. Cells expressing *CAT* mRNA with or without boxB were used. Levels of CAT activity in independent cell lines with inducible fusion protein expression were measured in the presence and absence of tetracycline. Lines without a lambdaN fusion were measured once with, and once without, tetracycline as a negative control. Expression of the tagged protein is shown in Supplementary Figure S7C and D. C) Migration of *CAT-boxB* mRNA in a sucrose gradient in the absence of λN-XAC1-myc. The membranes stained with methylene blue were used as a loading control. Quantification (bottom panel) was done for two independent experiments using ImageJ. D) Same as (C) but after induction of myc-XAC1-λN expression for 24 hours.

To check for effects of XAC1 tethering on translation we analysed the effect on migration of the *CAT-boxB* in a sucrose gradient. There was no reproducible effect of XAC1 over-expression on the total polysome profile, and total protein synthesis was not affected after 24h *XAC1* RNAi or XAC1 tethering (Supplementary Figure S1C). In the presence of XAC1, there was also no shift in *CAT-boxB* mRNA that might suggest an increase in ribosome density (Figure 7C, D). We previously found that MKT1 and PBP1 partially co-migrate with polysomes after sucrose gradient centrifugation (25). In contrast and rather surprisingly, most XAC1 appeared not to be polysome-associated (Supplementary Figure S8). This suggests either that many XAC1-PBP1-MKT1 complexes are not mRNA-bound or, less likely, that XAC1 was removed during centrifugation.

Finally we checked the effects of tethering some of the other proteins associated with XAC1 in bloodstream forms. First, full-length EIF4E6 was a clear activator (Figure 8A), which is consistent with the known activation by its partner EIF4G5 (46). We also tested two associated proteins of unknown function: we could not obtain full-length plasmid clones for Tb927.8.1290, but activation by Tb927.10.15310 (just below the significance cut- off, but present in all three XAC1 preparations), was confirmed (Figure 8B).

**Figure 8.**
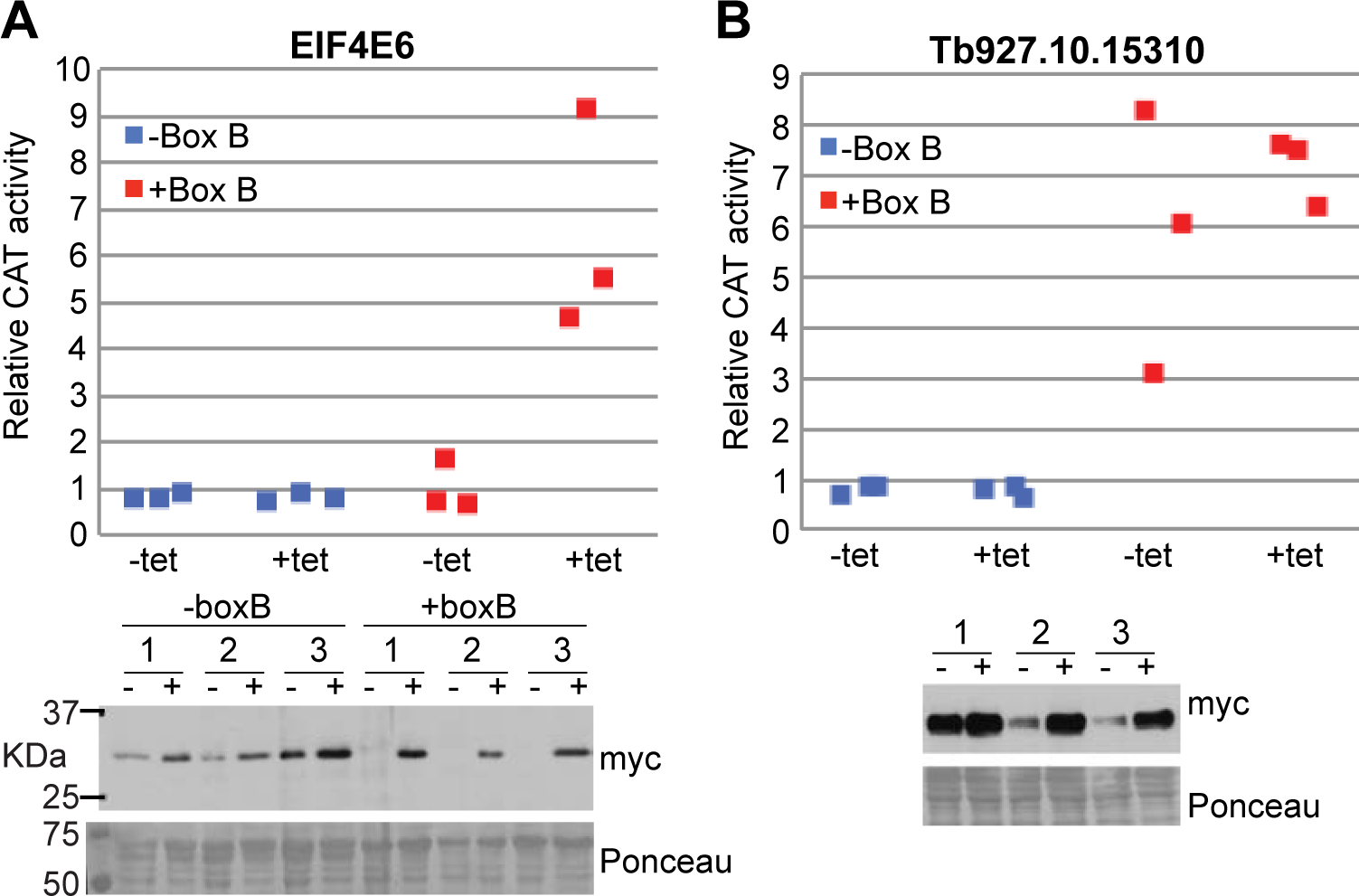
Activation by other tethered proteins. A) Tethering assay using EIF4E6. Cell lines expressing a *CAT* mRNA with or without five copies of boxB were used. Levels of CAT activity and *CAT* mRNA were measured in the presence and absence of tetracycline to induce lambdaN-EIF4E6-myc expression. B) Tethering assay using Tb927.10.15310. Details are as in (A).

### RNAs bound to MKT1-TAP and MKT1L-TAP

MKT1 and MKT1L should only enhance expression if they are associated with mRNA. In bloodstream forms, both MKT1L and MKT1 were found to be bound directly to poly(A)+ mRNA, although the result for MKT1 (false discovery rate 0.003) was more convincing than for MKT1L (false discovery rate 0.014) (46). We expect MKT1 to bind to mRNA via its interactions with RNA-binding proteins, but evidence that MKT1L has similar interactions was absent. To identify bound mRNAs, we therefore pulled down tandem affinity tagged proteins from lysates, released them by tobacco etch virus protease cleavage, and sequenced the associated RNAs.Reads were mapped to the *T. brucei* Lister 427 and TREU927 genomes. Results are in Supplementary Table S3.

For MKT1L, the yields of RNA in the pull-downs were low, so that multiple preparations had to be pooled. In one set of replicates, 66% of reads aligned to the Lister427 genome, but for the second replica only 9% did, and re-alignment to the human genome suggested that the majority of reads were human contaminants, which had presumably accumulated during the numerous purifications. Just 62 unique mRNAs were more than 2-fod enriched in both sample pools, with no enrichment for any particular category. Given the very low mRNA yields we concluded that MKT1L-TAP has only weak association with mRNAs.

For MKT1, bound mRNA was readily obtained, but principal component analysis did not clearly separate the bound and unbound fractions (Supplementary Figure S9A). The results overall suggested very broad mRNA binding by MKT1 (Supplementary Table S3). There was no overall enrichment of any functional class (Supplementary Figure S8B), or of expression site sequences, including the expressed variant surface glycoprotein, VSG222 (Supplementary Table S3, sheet 1). Notably, however, mRNAs encoding ribosomal proteins were significantly less MKT1-bound than other mRNAs (Figure 9A, Supplementary Figure S9B).

**Figure 9.**
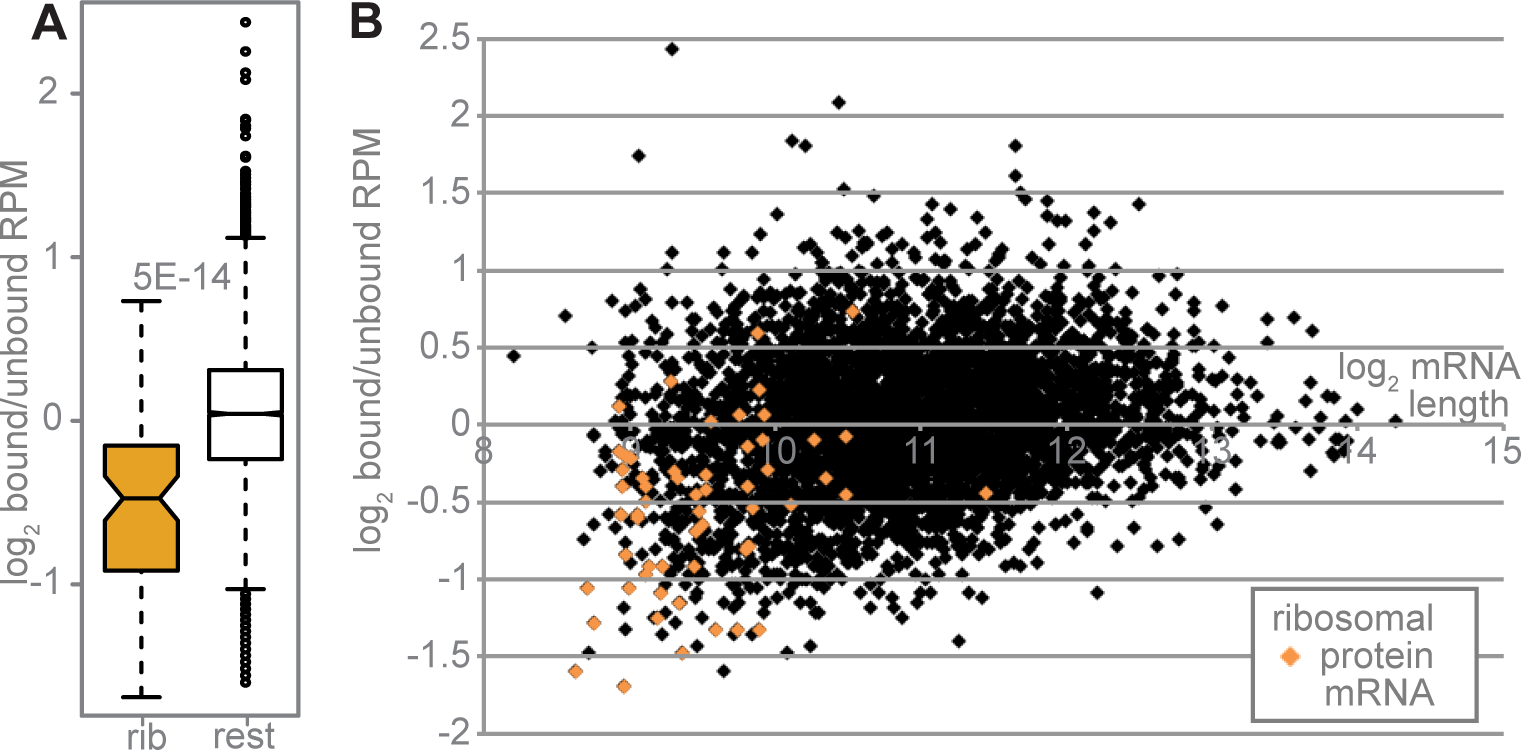
mRNAs encoding ribosomal proteins show low association with MKT1. The RPM of each unique mRNA in the MKT1-TAP bound eluate was divided by the RPM in the unbound fraction. The averages of the two values, log2-converted, were plotted. A) Comparison of mRNAs encoding ribosomal proteins with all other mRNAs. The boxes extend to the 25th and 75th percentiles. The whiskers extend to the most extreme data point which is no more than 1.5 times the interquartile range from the box. “rib” - mRNAs encoding ribosomal proteins. “rest” - all other mRNAs. An analysis for all functional classes is shown in Supplementary Figure S8. B) Relationship between binding fraction (bound/unbound) and annotated mRNA length. Results for mRNAs encoding ribosomal proteins are in orange.

These mRNAs are unusually short, but overall, there was no correlation between average MKT1 binding and the mRNA length (Figure 9B), which argues against completely non-specific binding. There was also no correlation with coding-region or 3’-UTR length, or with ribosome density (Supplementary Figure S9C). Overall these results indicate that MKT1 is associated with a large number of mRNAs, consistent with its interactions with many different RNA-binding proteins. The only associated proteins for which the mRNA specificities are known are ZC3H11 (17), ZC3H20 (18) and PUF9 (55). No enrichment of their targets was found, but ZC3H11 and ZC3H20 have low abundances in bloodstream forms, and PUF9 acts only in S phase.

## Discussion

Our results show that trypanosome PBP1, XAC1 and LSM12 form at least two different complexes. One contains MKT1, and the other contains MKT1L. The results of two-hybrid analysis indicate that MKT1L, like MKT1, joins the complex via PBP1. In bloodstream forms, the MKT1-containing complex is associated with numerous different RNA-binding proteins and mRNAs. In contrast, we found no convincing evidence for interaction of MKT1L with either RNA or RNA-binding proteins.

Although MKT1L is essential, our results yielded no insight into its precise function. We found no clear association with mRNA and no bound RNA-binding proteins, but this has to be viewed with caution because C-terminally TAP-tagged protein appeared to be only partially functional. Results from co-purifications and native gel electrophoresis suggested that most MKT1L is present as homo-dimerized PBP1-LSM12-XAC- MKT1L tetramers; very limited association between the two different complexes might account for occasional co-purification of MKT1 and MKT1L. This association could occur through homo-dimerization of either PBP1 or PABP. We considered the possibility that MKT1L competes with MKT1 for binding to PBP1, giving a complex that is inactive in RNA metabolism and thus regulating MKT1 action. Another possibility is that MKT1L helps to stabilise PBP1, maintaining a “store” for future MKT1 interactions. Intriguingly, a version of MKT1L that lacked the N-terminal extension interacted with EIF4E6, whereas neither full-length MKT1L, nor MKT1, did so. It is therefore possible that when MKT1L is present in complexes that include MKT1, it helps to stabilize interactions with the EIF4E6/G5 complex. This activity might be regulated by the N-terminal extension.

Messenger RNA co-purified with MKT1, but relatively few individual mRNAs were strongly enriched. We therefore suggest that association of MKT1-containing complexes with mRNAs is relatively promiscuous, with the rather mild preferences for some mRNAs over others being guided by a spectrum of cooperative interactions, not only with specific RNA-binding proteins, but also (via PBP1) with poly(A)-tail bound PABP2 and perhaps also with the EIF4E6/4G5 initiation complex (see below). Although we expected that MKT1- bound mRNAs would be stable and well-translated, there was in fact no correlation with mRNA half-life (13) (not shown), ribosome density (Supplementary Figure S9C) (12) or the procyclic form codon optimality index (geCAI from (8) (not shown). The easiest explanation for this is that several factors influence the behaviours of mRNAs, including codon optimality as well as a spectrum of associated proteins. The action of MKT1 complexes may well compete with destabilising or repressive proteins that are bound to other parts of the mRNA.

Notably, the mRNAs encoding ribosomal proteins were relatively depleted in the MKT1-bound fraction. These mRNAs are notable for high codon optimality than other mRNAs (8) (Supplementary Figure S10), which might explain their rather low ribosome densities (12). They also have unusually short untranslated regions (15) so may not associate with many RNA-binding proteins.

*T. brucei* DHH1, a homologue of *S. cerevisiae* Dhh1 and metazoan DDX6/Rck, co-purified with XAC1 (this paper), MKT1 and ZC3H11 (25). Numerous, mostly suppressive, roles have also been reported for the Opisthokont DHH1 homologues, including decapping activation (56) and promoting deadenylation (57). The Apicomplexan version of DHH1, DOZI, is also implicated in translation repression (58). Kinetoplastid decapping is completely different from decapping in Opisthokonts (59), but DHH1 is indeed associated with the *T. brucei* CAF1-NOT complex (60). *T. brucei* DHH1 is, correspondingly, required for developmental down- regulation of mRNAs, but expression of a mutant version resulted in overall translation inhibition, suggesting more generalised effects. Moreover *T. brucei* DHH1 has appeared in many other mRNA-related purifications, including splicing factors, suggesting a variety of roles in mRNA metabolism (45,61,62). We do not know what, if any, role DHH1 plays in MKT1-complex function.

MKT1 was previously shown to interact with several proteins that have known RNA-binding domains, and/or can be cross-linked to mRNA (46). Our affinity purification of XAC1 provides more robust data concerning *in vivo* association of these proteins, and confirms the association between interaction with the MKT complex and activation in the tethering assay (Table 1) (46, 47). ZC3H20 is more abundant in stumpy-form and procyclic trypanosomes than in bloodstream forms (18, 48), but was nevertheless detected in the bloodstream-form XAC1 preparation. Its mRNA targets are enriched for procyclic-form-specific surface proteins and mitochondrial components (18, 19). PUF9 stabilises some S-phase mRNAs (55). Nothing is known about the functions of ZC3H28, PUF6 and the associated proteins of unknown function, beyond their cytoplasmic location (54), mRNA binding and activating potential. ZC3H32 suppresses expression when tethered but no clear mRNA targets were identified (44). CFB2 is quite intriguing: it is an essential mRNP protein (46, 63) with an MKT1 interaction motif; but it also has an F-box domain, which may explain the presence of SKP1 (which interacts with F-boxes) in the XAC1 pull-down. Additional RNA-binding proteins that were enriched with XAC1, but below the significance cut-off, were SLBP1, ZC3H5, ZC3H34, UBP2, ALBA3, and the known MKT1 partner ZC3H11, which is very difficult to detect by mass spectrometry (64).

XAC1-associated RNA-binding or mRNP proteins that were not previously found with MKT1 included DRBD2 and DRBD3/PTB1, both of which have a polyglutamine repeats. DRBD2 is an orthologue of yeast Gbp2, which is implicated in mRNA export and quality control. Neither MKT1 nor PBP1 was found by mass spectrometry of *T. cruzi* DRBD2 preparations (65) and it repressed in the tethering screens. DRBD3/PTB1 has roles in both mRNA stablisation (66–68) and processing (68), targeting preferentially mRNAs whose encoded proteins have roles in translation and proline metabolism (69).

Although PBP1 interacts with both PABP1 and PABP2 in the yeast 2-hybrid assay (25), only PABP2 co-purified with XAC1. This result is consistent with a published study: PBP1 was enriched in pull-downs of PABP2, but not PABP1 (70). Procyclic-form PABP2 pull-downs, especially under low-salt conditions, also contained the MKT1 complex and several MKT1-interacting proteins, including ZC3H20, ZC3H28, ZC3H39, PUF6 and PUF9.

One of the most interesting results was the preferential association of XAC1-containing complexes with just one of the five possible EIF4E-EIF4G translation initiation complexes, containing EIF4E6 and EIF4G5.

Moreover, EIF4G5 specifically interacts with MKT1 in the 2-hybrid assay. G5-IP, which interacts with EIF4G5 (49), was also detected in two out of three XAC1 preparations, but not the control. The EIF4E6/G5 complex preferentially associates with PABP2 (49, 70), just like XAC1. Depletion of EIF4E6 in procyclic forms results in viable cells with detached flagella (49), and both EIF4E6 (this paper) and EIF4G5 (46) activate when tethered, which suggests that this complex is active in translation, although its specificity is unknown. Indeed, the translation helicase eIF4A1 was in the XAC1 purification and was not detected with either GFP or MKT1L. We therefore suggest that MKT1 complexes can enhance translation not only through binding to PBP1-PABP2, but also by direct recruitment of EIF4E6/4G5.

## Experimental procedures

### Cells and plasmids

Details of all plasmids and oligonucleotides are provided in Table S1. The culture conditions were as described in (71). All experiments were done with Lister 427 monomorphic bloodstream parasites expressing the Tet-repressor and with procyclic forms expressing the Tet-repressor and T7 polymerase (72).

Stable cell lines expressing V5- or YFP- tagged proteins were generated by the integrating the tag-encoding sequence in frame with the coding region at the original locus (73, 74). MYC- and TAP- tagged proteins were obtained by tetracycline-inducible expression of fusion proteins from constructs integrated in the rRNA locus (75). Expression was induced using 200ng/ml tetracycline. RNAi against XAC1 or MKT1L was done using inducible stem-loop plasmids in cell line with one of the copies of XAC1 or MKT1L tagged with V5.

For the tethering assays, cell lines constitutively expressing CAT reporter with actin 3′-UTR or boxB actin 3′- UTR were co-transfected with plasmids for inducible expression of inducible λN-XAC1-myc full-length or fragments fusion proteins (41).

For gene knockout assays, deletion cassettes were constructed using the Blasticidin S deaminase (*BSD*), Puromycin acetyltransferase (*PAC*) or Neomycin phosphotransferase (*NPT*) genes. After transfection and selection the desired resistance marker replaced one or both alleles of MKT1 or MKT1L.

### 35S Pulse Labeling

3×10^7^ cells were spun down at 800 g for 10 min at RT, transferred to a 1.5 ml Eppendorf tube and washed twice with 1x PBS followed by centrifugation at 4,000 g for 3 min. The cell pellet was resuspended in 400 µl labeling medium and incubated at 37°C for 15 min. 2 μl of L-[^35^S]-methionine with an activity of about 20 μCi, was added. The cells were incubated for 1 h at 37°C. The cells were pelleted and washed twice with 1x PBS wash buffer (4,000 g for 3 min at RT). The pellet was resuspended in 15 μl of Laemmili lysis buffer and the proteins were separated in a 12% SDS gel.

The labelling medium was Dulbecco’s modified eagle medium (GIBCO, high glucose, containing pyridoxine hydrochloride, lacking L-glutamine, sodium pyruvate, L-methionine and L-cysteine), supplemented with 25 mM HEPES, 2 mM glutamine, 0.1 mM hypoxanthine, 1.5 mM L-cysteine, 0.0028% β-mercaptoethanol, 0.05 mM bathocupronsulfate and 10% heat-inactivated FCS (previously dialyzed against 30 mM HEPES pH 7.3 / 150 mM NaCl).

### Targeted yeast two-hybrid analysis

The Matchmaker Yeast Two-Hybrid System (Clontech) was used for protein–protein interaction analysis following the manufacturer’s protocol. The DNAs of the protein ORFs were PCR-amplified and cloned into both pGBKT7 and pGADT7. The plasmids were co-transformed pairwise into AH109 yeast strains (Matchmaker 3 System, Clontech), and selected initially on double drop-out (DDO) medium (minimal SD media lacking Trp and Leu) on quadruple drop-out (QDO) medium (minimal SD media lacking Trp, Leu, His and Ade). Positive interactions were indicated mainly by growth on quadruple drop-out (QDO) medium (minimal SD media lacking Trp, Leu, His and Ade) and sometimes confirmed by a blue colour change due to X-α-gal present in the medium. The interaction between murine p53 and SV40 large T-antigen served as positive control, with LaminC and SV40 large T-antigen as negative bait (DNA-binding domain) and prey controls.

### Affinity purification for mass spectrometry

For tandem affinity purification from bloodstream forms, approximately 1×10^10^ trypanosomes expressing TAP-tagged protein were harvested and used for tandem affinity purification as previously described (76). The elute was run 2 cm into a 1.5 mm NuPAGE™ Novex™ 4-12% Bis-Tris protein gel (Thermo Fisher Scientific), stained with Coomassie blue and distained with distaining solution (10% acetic acid and 50% methanol). The protein-containing gel area was analysed by mass spectrometry. Data were analysed quantitatively. Affinity purifications from procyclic forms were done using cells expressing V5 *in situ* tagged proteins. 1x 10^9^ cells were harvested by centrifugation at 2300g for 13 min, 4°C. The cell pellet was washed in 10 ml of ice cold PBS supplemented with protease inhibitors, followed by centrifugation at 2.300g, 8 min. 4°C. Cell lysis was done by adding 1 ml of IP lysis/wash buffer with low salt concentration (20mM Tris pH 7.5, 150mM KCl, 5 mM MgCl_2_, 0.05% IGEPAL, 0,5 mM DTT) supplemented with 10 ug/ml of aprotonin and 10 ug/ml of leupeptin. Cells were pipetted up and down, passed 20 times through a 21G x 1^1/2^ needle and 20 times through a 27G x ¾ needle.

The complete cell lysis was checked under light microscope slides. The sample was centrifuged at 10.000g, 15 min, 4°C and the supernatant incubated during 4 h at 4°C with 50 µl of V5 magnetic beads (20 mg/mL of Dynabeads® M-280 Tosylactivated, Invitrogen, coupled with Anti V5-Tag Antibody, clone SV5-Pk1, Bio-Rad). The unbound fraction was discarded and the beads washed 2 times with 1ml of IP Buffer and additionally twice more with 1 ml ice cold PBS + protease inhibitors. The bound proteins were neutrally eluted with 50 µL of V5 peptide (V7754-4MG, SIGMA) at 1mg/mL in PBS, at 37**⁰**C on a rotator for 10 min. The eluate sample was submitted to mass spectrometry as described for the bloodstream forms.

### Quantitative Mass spectrometry data analysis

Nanoflow LC-MS^2^ analysis was performed at the ZMBH mass spectrometry facility, with an Ultimate 3000 liquid chromatography system directly coupled to an Orbitrap Elite mass spectrometer (both Thermo-Fischer, Bremen, Germany). Samples were delivered to an in-house packed analytical column (inner diameter 75 µm x 20 cm; CS – Chromatographie Service GmbH, Langerwehe, Germany) filled with 1.9 µm ReprosilPur-AQ 120 C18 material (Dr. Maisch, Ammerbuch-Entringen, Germany). The mass spectrometer was operated in data-dependent acquisition mode, automatically switching between MS and MS^2^. For data analyses, the MaxQuant platform (version 1.5.3.30) was used for peak list picking, protein identification and validation. Protein identification was based on the *T. brucei brucei* 927 annotated protein database, release 9.0 from TriTrypDB. For validation, a minimum of seven amino acids for peptide length and two peptides per protein were required. In addition, a false discovery rate (FDR) threshold of 0.01 was applied at both peptide and protein levels. Peptides shared between orthologues were assigned to the same protein group.

Statistical analysis of the results was performed using the Perseus toolbox (version 1.6.2.1). Ratios were calculated from label-free quantification intensities. Missing values were imputed using the Perseus default settings of imputation (mean down shift = 0.3, standard deviation down shift = 1.8). A two sample t-test was performed comparing the natural logarithm of the intensities from the experiments and the control groups. To confirm the specificities of the purifications, data from TAP-GFP or V5-GFP purifications were used as controls for bloodstream- and procyclic-form purifications, respectively.

### Affinity purification and RNASeq

For RNAseq analysis, we used cells expressing TAP-tagged protein but omitted the second step of purification. The bound and unbound RNAs were purified using peqGOLD Trifast reagent (Peqlab), and rRNAs were depleted using rRNA-complementary oligonucleotides and RNase H (77). Library-building, sequencing and the data analysis were done as described in (78).

Data were trimmed and aligned using the tryprnaseq (79) pipeline, which includes Bowtie2. Each read was allowed to align once to both the Lister427_2018 (80) and TREU 927 (81) genomes. The Lister 427 genome corresponds to the cells used for the experiments, and includes strain-specific expression sites and Variant Surface Glycoprotein genes. However, the TREU927 genome is much better annotated in chromosome-internal regions. When genes are present more than once, which is common in trypanosomes, the reads are shared among the different copies. The ploidy of each of the “unique” genes was assessed using two datasets, for which each read was allowed to align twenty times.

To look at enriched gene categories, we analysed the data only for one representative of each repeated TREU927 gene using a list modified from (9) (see Supplementary Table S3). To ensure that each sequence was weighted appropriately, before the calculations, each “unique gene” read count was multiplied by the ploidy. Data were analysed using DeSeqUI (82), an adapted version of DeSeq2 (83).

ArrayExpress accession numbers are as follows: XAC1 yeast 2-hybrid screen: E-MTAB-4946; XAC1 RNAi: E-MTAB-6239; TAP-MKT1 pull-down: E-MTAB-6907; TAP-MKT1L pull-down: E-MTAB-6904.

### Northern blotting and polysome analysis

For Northern blotting, total RNA was extracted using peqGOLD Trifast reagent (Peqlab). Isolated RNA (typically 20µg of total RNA or RNA purified from entire gradient fraction) was resolved on formaldehyde agarose gel, blotted to nylon membranes and detected by hybridization with radioactive probes for *CAT*, and β-tubulin (Tb927.1.2370) mRNAs. Quantification was done using ImageJ Software.

Polysome fractionation was done using 5×10^8^ bloodstream-form cells (1×10^6^ cells/ml) as described in (77). Total RNA was extracted from each fraction as above and examined by Northern blotting.

### Protein detection and manipulation

Immunoprecipitation assays were done as previously described (20). 1×10^8^ bloodstream trypanosomes expressing the tagged protein were harvested by centrifugation (800 g, 10 min, 20°C), washed with 1 ml of cold phosphate-buffered saline and lysed in hypotonic buffer (10 mM NaCl, 25 mM Tris-Cl pH 7.5, 10 µg/ml leupeptin, 0.1% IGEPAL) by passing 10 times through a 21G needle. After pelleting insoluble debris by centrifugation (17000 g, 10 min, 4°C) and adjusting to 150 mM NaCl, the clarified lysate was used for immunoprecipitation with 30 µl of anti-myc or anti-V5 coupled beads (Bethyl Laboratories). The YFP-XAC1 fusion protein was immunoprecipitated using GFP binding protein coupled to NHS-Sepharose that were kindly given by Dr. Georg Stoecklin, ZMBH, Heidelberg University, Germany. The beads were incubated for 2 hours at 4°C and then washed 4 times at 4°C (5 min of incubation follow by centrifugation at 850 g, 5 min) with IPP150 (10 mM Tris pH 7.5, 150 mM NaCl, 0.1% IGEPAL). Samples for western blots were taken during the procedure and the beads were boiled in SDS sample buffer. Proteins were detected by western blotting according to standard protocols. Chloramphenicol acetyltransferase activity was measured in a kinetic assay involving partition of ^14^C-buturyl chloramphenicol from the aqueous to the organic phase of scintillation fluid (84). The total protein concentration was measured by the Bradford method.

### Blue Native Gel

Samples for Native gel analysis were prepared as follow: 1×10^8^ bloodstream trypanosomes expressing the *in situ* N terminal V5 tagged protein were harvested by centrifugation at 800 g, 8 min at room temperature, washed twice with 1 ml of cold 1X PBS, resupended in 50 µl of Native extraction buffer (25 mM HEPES, 150 mM sucrose, 20 mM potassium glutamate, 3 mM MgCl_2_,0,5% Igepal, 150 mM KCl, 0,5 mM DTT, 10 µg/ml leupeptin, 10 µg/ml aprotonin), incubated on ice for 30 minutes, and spun down at 17000 g for 15 min at 4°C. Supernatants were prepare and electrophoresed in a precast 4-16% NativePAGE, Novex Bis-Tris Mini Gel following manufacturer’s protocol (Life technologies). Native Mark Unstained Protein Ladder (Life technologies) was used as standard protein sizes. Proteins were transferred to 0.45µm Immun-Blot PVDF membranes (Thermo Scientific™ Pierce™) with 1X NuPAGE® Transfer Buffer at 25V, 4°C, overnight.

Membranes were fixed in 8% acetic acid for 15 min, rinsed with water, followed by regular Western Blot procedure with specific antibody.

## Acknowledgements

This work was supported by grant number Cl112/17-2 to Christine Clayton, and by core support from the State of Baden-Württemberg. Bin Liu was partially supported by a Humboldt stipendium and Franziska Egler had a stipend from the Heidelberg Biosciences graduate school (HBIGS).

We thank Kwado Oworae for cloning, pHD 3097,3096,3095 and Alexandra Dias for transfections and selections of parasite with those constructions; Dr. Esteban Erben for the cell line of bloodstream forms expressing *in situ* V5-XAC and Marvin Gathof for contributions to Y2H experiments. The Core Facility for Mass Spectrometry & Proteomics (CFMP) at the ZMBH and the Deep Sequencing core facility at the BioQuant.

## Author contributions

L.N and M.T. did the RNAi experiments, in procyclic and bloodstream forms, respectively (Figure 1 and Figure 5). L.N and M.T. performed Protein purifications and Mass Spectrometry experiments and L.N analysed the Proteomic data (Figure 2, Table 1 and Supplementary Table S2). L.N. and M.T. did RNA pull-down and sequencing and K.M. and C.C. analysed the RNA pull down data (Figure 9 and Supplementary Table S3). L.N did protein quantification and Native complex composition experiments (Figure 3), Y2H experiments (Figure 4A, B and Supplementary Figure S4 and S5) and MKT1/MKT1L knockouts (Figure S6). F.E did the co- immunpreciptations of EIF4E6 and MKT1/ MKT1L and B.L co-immunopreciptations of ZC3H20/21 and MKT1/MKT1L (Figure 4C, D and Supplementary Figure S2). M.T and KM did the immunofluorescence experiments (Figure 6) and tethering experiments (Figure 7, 8 and Supplementary Figure S7). M.T. was responsible for polysome purifications and Northern Blots (Supplementary Figure S1C and S9). C.C. conducted data analysis, funding acquisition, writing and editing manuscript. We thank Claudia Hezmbh.,lbig for technical assistance.

---

**Figure S1.**
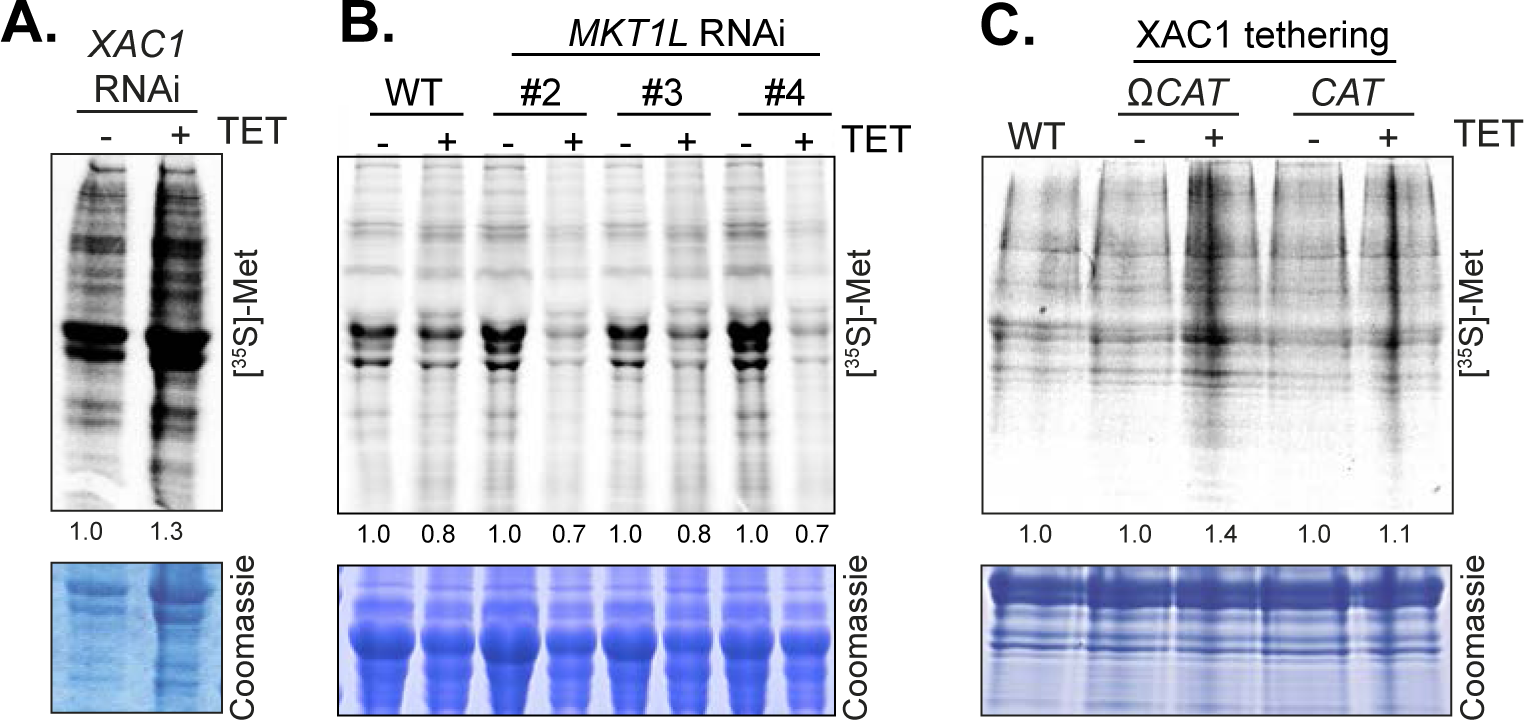
Influence of XAC1 and MKT1L changes on translation. Cells were pulse labelled for 30 min with [^35^S]-Methionine and the resulting extracts were analysed by SDS-PAGE and autoradiography. Assays were done 24h after tetracycline addition. A) Cells with *XAC1* RNAi B) *MKT1L* RNAi. C) Inducible expression of lambdaN-XAC1

**Figure S2.**
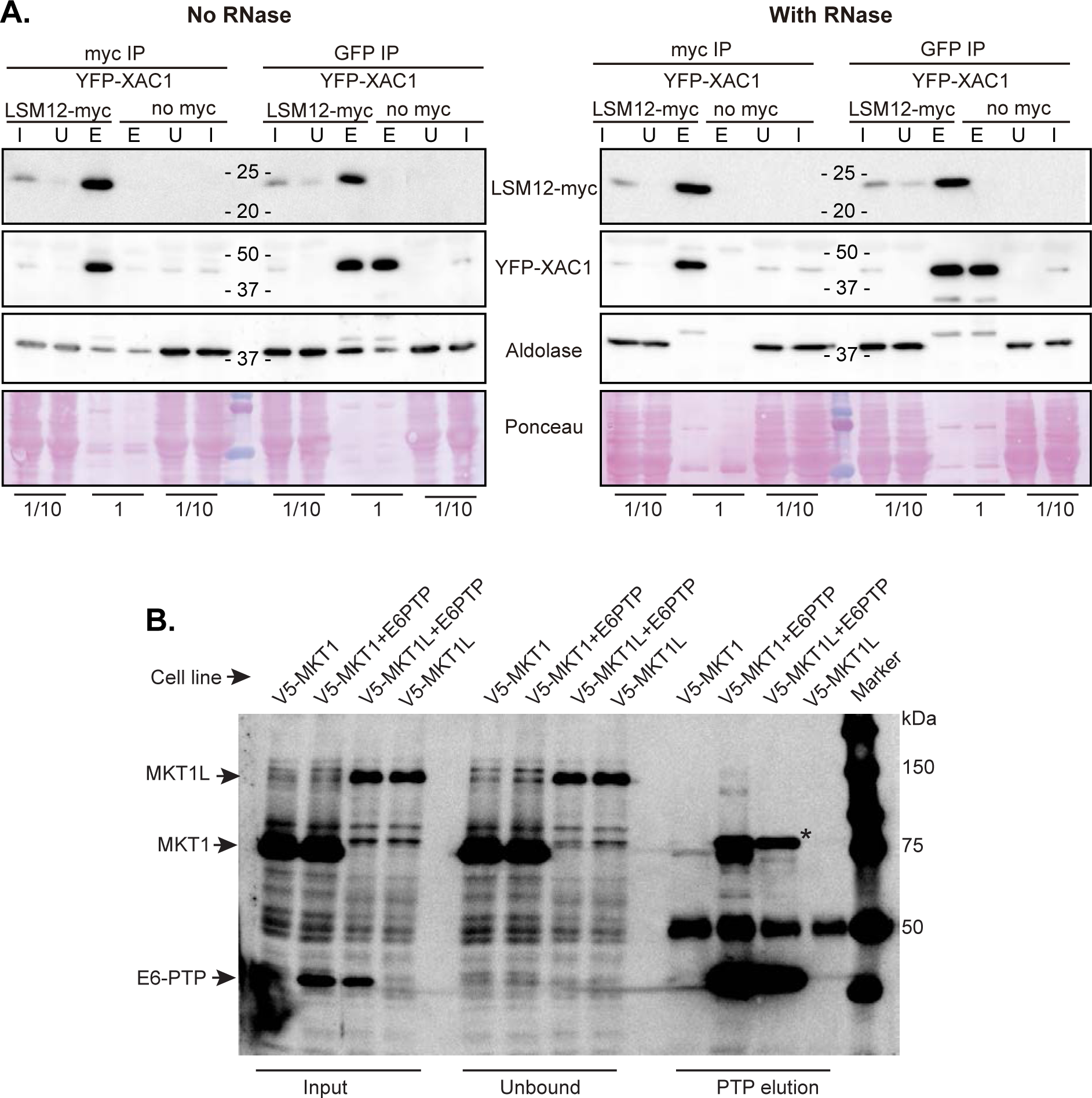
XAC1-LSM12 interaction and EIF4E6-MKT1/MKT1L interactions. A) Mutual immunoprecipitation of YFP-XAC1 (*in situ* tagged) and inducibly expressed LSM12-myc. Aldolase and the Ponceau staining were used as loading controls. I: input, 1/10 of the total cell extract, U: unbound fraction - same volume as of the input fraction; E: all of the eluate. B) Over-exposed version of Figure 4C.

**Figure S3. Sequence alignments for MKT1 and MKT1L** The alignment was created using Clustal Omega (EMBL-EBI). Organism suffixes are: *Trypanosoma cruzi -* TcCLB; *Trypanosoma vivax -* TvY; *T. brucei -* Tb; *Trypanosoma congolense -* TcIL; *Blechomonas ayalai* - Baya; *Crithidia fasciculata -* CFAC1; *Leishmania major -* LmjF; *Endotrypanum monterogeii* - EMOLV. The colour key is as in Figure 1. The PIN domain is grey, the MKT-N domain is rgeen and the MKT-C domain is cyan. Arrows are as follows: (1) D->G in S. cerevisiae Mkt1 affects gene expression, gives better growth at 41°C and better sporulation. (2) S288C S. cerevisiae has D->A. This inactivates Mkt1. (3) This is D in XPD1 and Fen1 endonucleases. D->A is inactive, D->E is active. (4) This is E in XPD endonuclease. Mutation to A: can still bind DNA but is inactive. (5) A->V in XPD has reduced activity. (6) In XPD, D->A abolishes 3’ excision.

**Figure S4.**
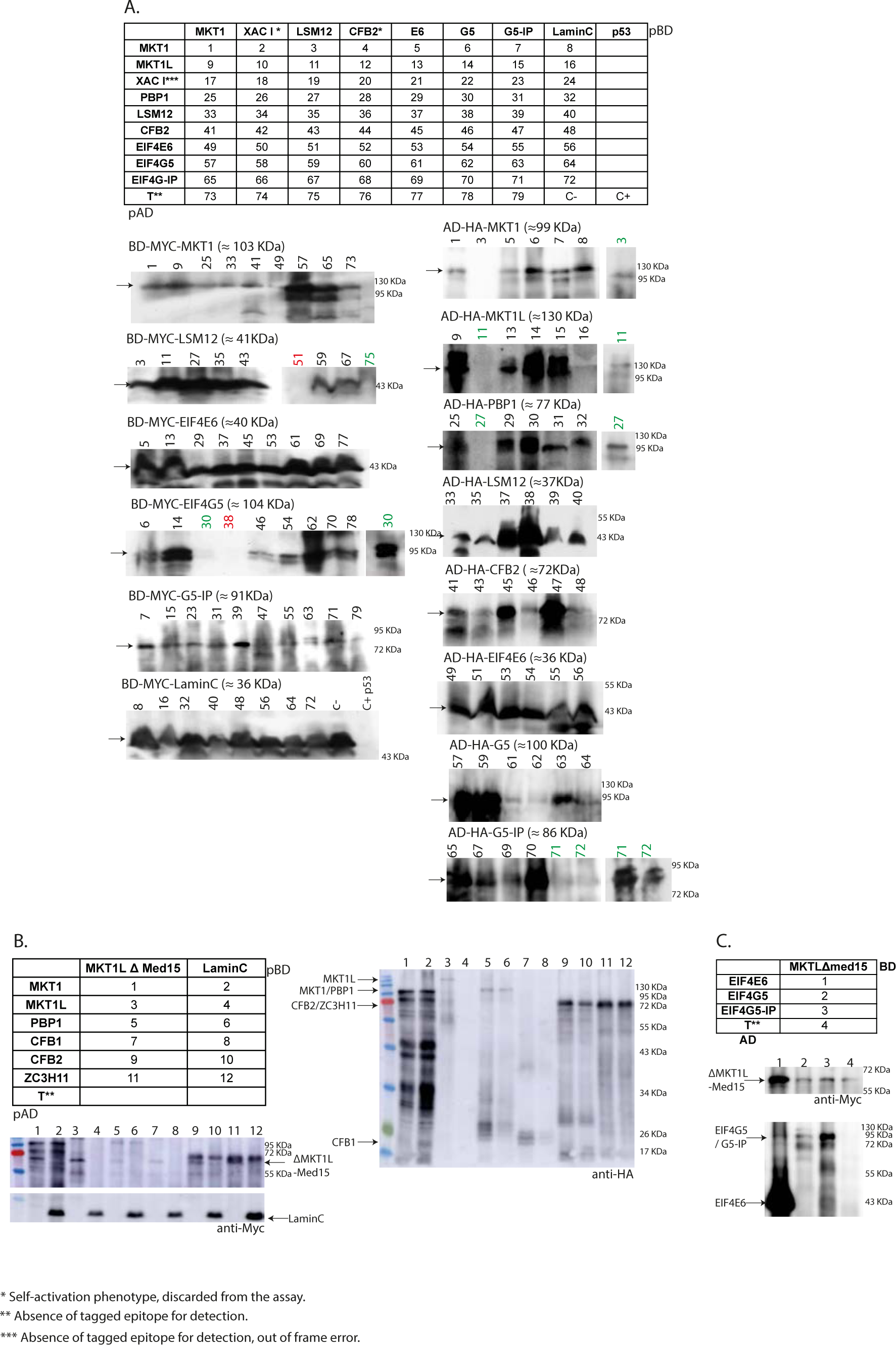
Expression of activation domain and binding domain hybrid proteins. A) AH109 yeast strains were transformed with the indicated plasmid combinations of bait (pBD) and prey (pAD) as described at upper table. Bellow, Western blot detection of fusion BD proteins (Myc tag) and AD proteins (HA tag). *, Self-activation phenotype, discarded from the assay. **, Absence of tagged epitope for detection. ***, Absence of tagged epitope for detection, out of frame error. Numbers in red, first WB detection failed. Numbers in green, WB repetitions. B) Detection of fusion proteins from additional interactions of MKT1LΔN and RBPs. Details as in A). C) Detection of fusion proteins from additional interactions of MKT1LΔN and EIF’s. Details as in A).

**Figure S5.**
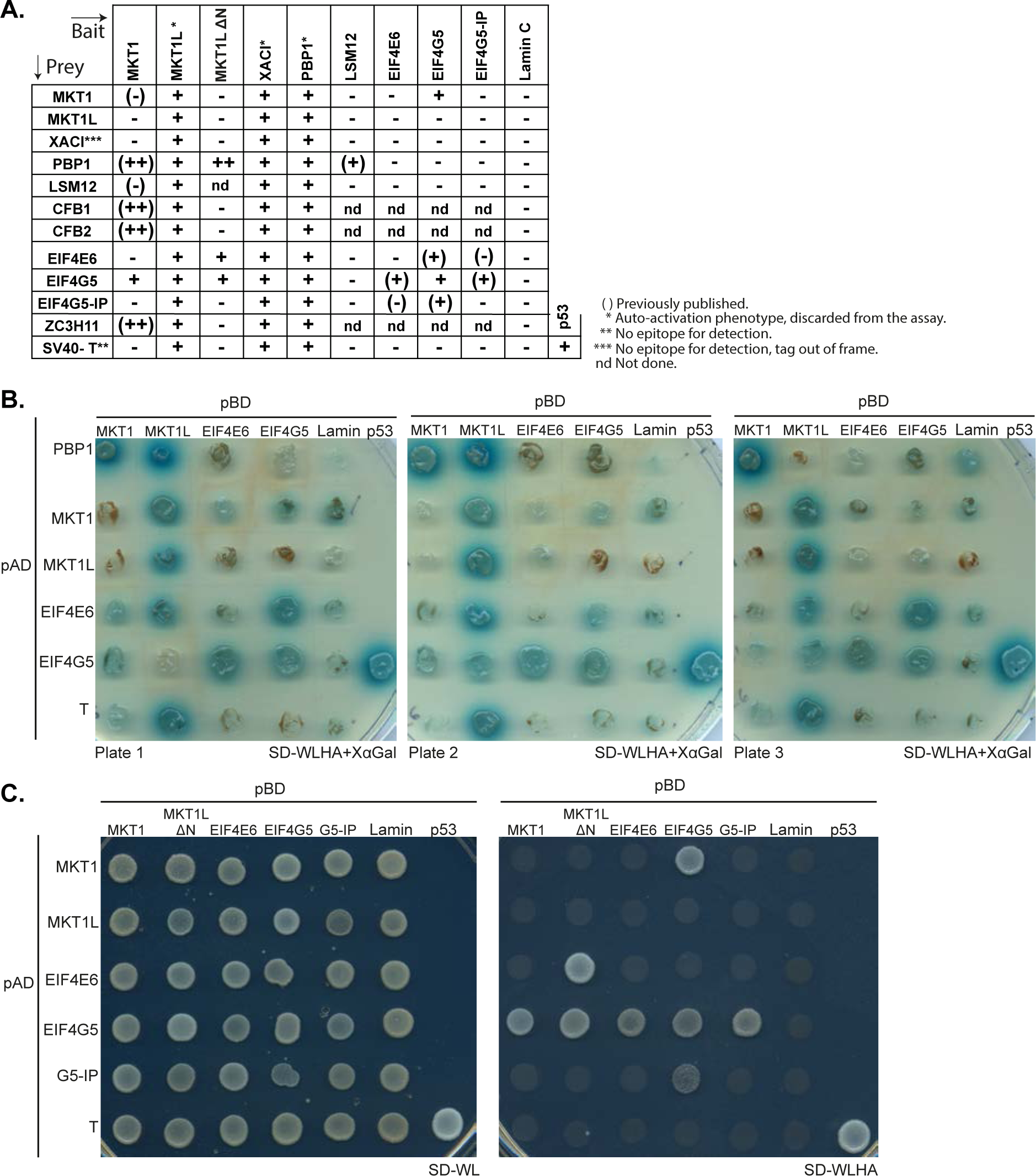
Additional 2-hybrid results. A) Summary table of all tested interactions. In parentheses, previously published interactions. *, Self-activation phenotype, discarded from the assay. **, Absence of tagged epitope for detection. ***, Absence of tagged epitope for detection, out of frame error. B) Individual pictures were pasted together to give an overview of the results. Transformed yeast cells were grown on selective media lacking Ade, His, Leu, Trp and plus X-α-Gal (-LTHA+ X-α-Gal) to test protein interaction (Plates 1 to 3). C) Transformed yeast cells were grown on selective media lacking Leu and Trp (-LT) as control (SD-WL panels), and on media lacking Ade, His, Leu, Trp and plus X-α-Gal (-LTHA) to test protein interaction (SD-WLHA panels).

**Figure S6.**
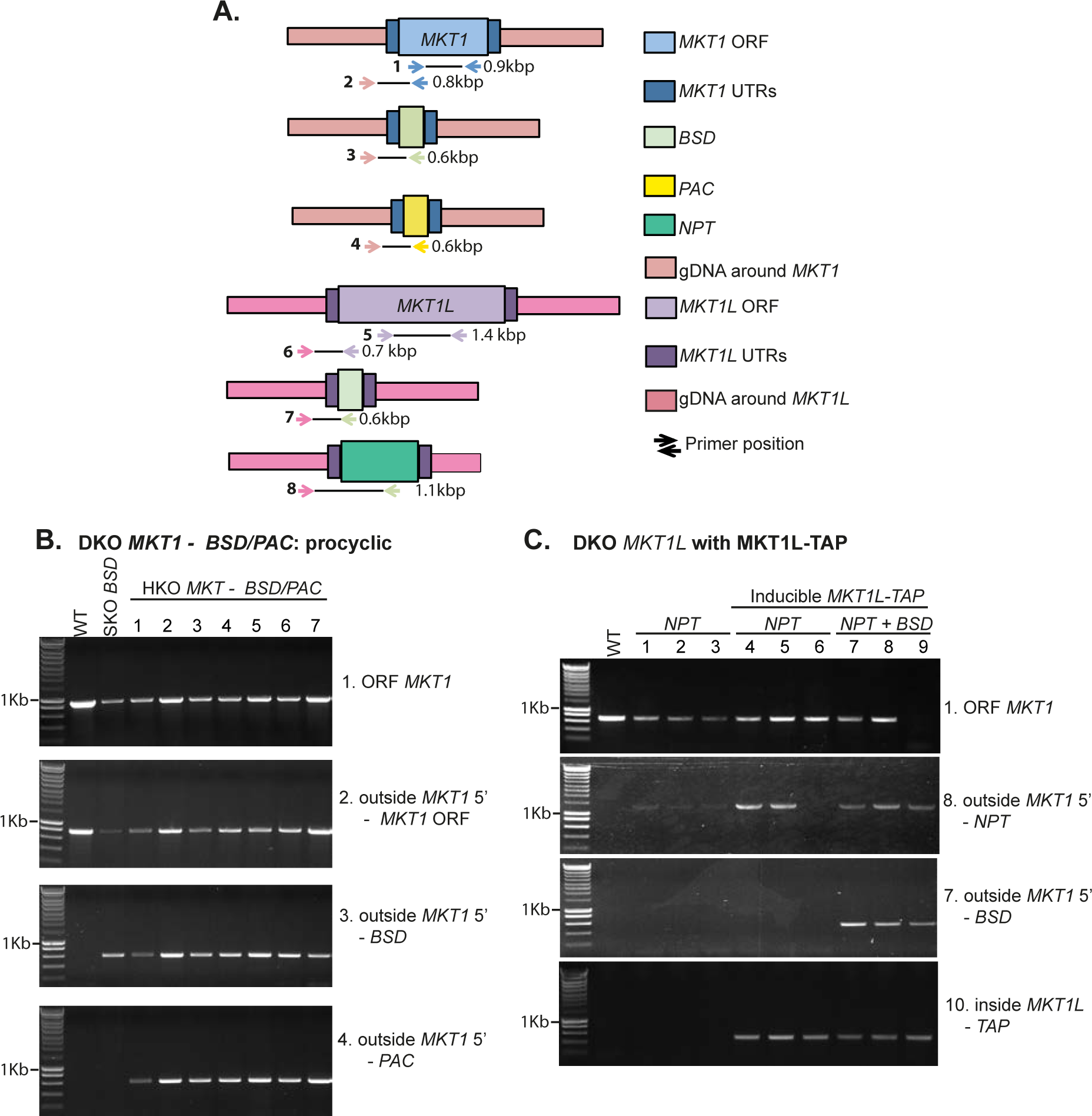
Examples of clone verification. A) Schematic maps of the MKT1 and MKT1L loci, and of plasmids used for gene replacement, along with oligonucleotide primers used for PCR. B) A line in which one *MKT1* gene had been replaced by the *BSD* gene (SKO-BSD) was transfected with a fragment designed to replace the second allele with *BSD*. PCRs were done using genomic DNA from individual clones and the primers shown in (A). C) Replacement *MKT1L* genes with *BSD* and *NPT*. Lanes 1-3 are independent clones in which one *MKT1L* gene was replaced with *NPT*. For lanes 4-9, cells inducibly expressing MKT1L-TAP were used. First, one endogenous copy of *MKT1L* was replaced with *NPT*. Then, tetracycline was added (100 ng/ml) and the cells were transfected with a construct designed to replace the second endogenous copy of *MKT1L* with the *PAC* resistance marker. The clone shown in lane 9 no longer has any copies of the endogenous constitutively-expressed gene. Those in lanes 7 and 8 have both resistance markers but have also retained the endogenous gene. The oligos used for amplification of the TAP-tagged gene copy are not illustrated.

**Figure S7.**
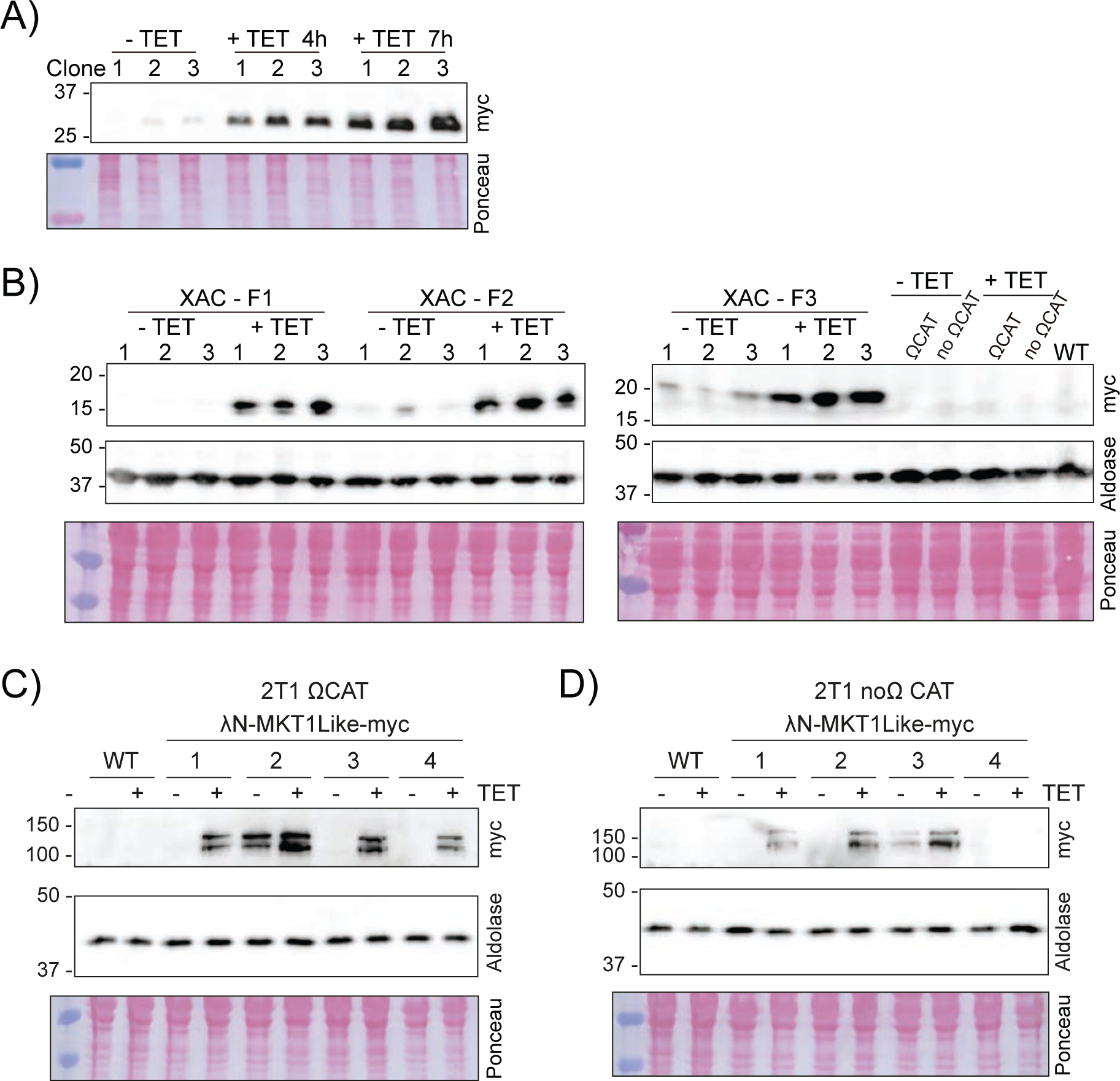
λN-protein-myc expression controls. A) λN-XAC1-myc, time course of expression. B) λN-XAC1-myc fragments expression. The cell line expressing the CAT with (ΩCAT) or without (no ΩCAT) the boxB sequence were used as controls. Expression was induced for 24h. C) λN-MKT1Like-myc expression in the cell line expressing the CAT with the boxB sequence. The clones 1, 3 and 4 were used in the tethering experiments. Expression was induced for 24h. D) λN-MKT1Like-myc expression in the cell line expressing the CAT without the boxB sequence. The clones 1, 2 and 4 were used in the tethering experiments. The aldolase gene or the Ponceau staining were used as a loading control. Expression was induced for 24h.

**Figure S8.**
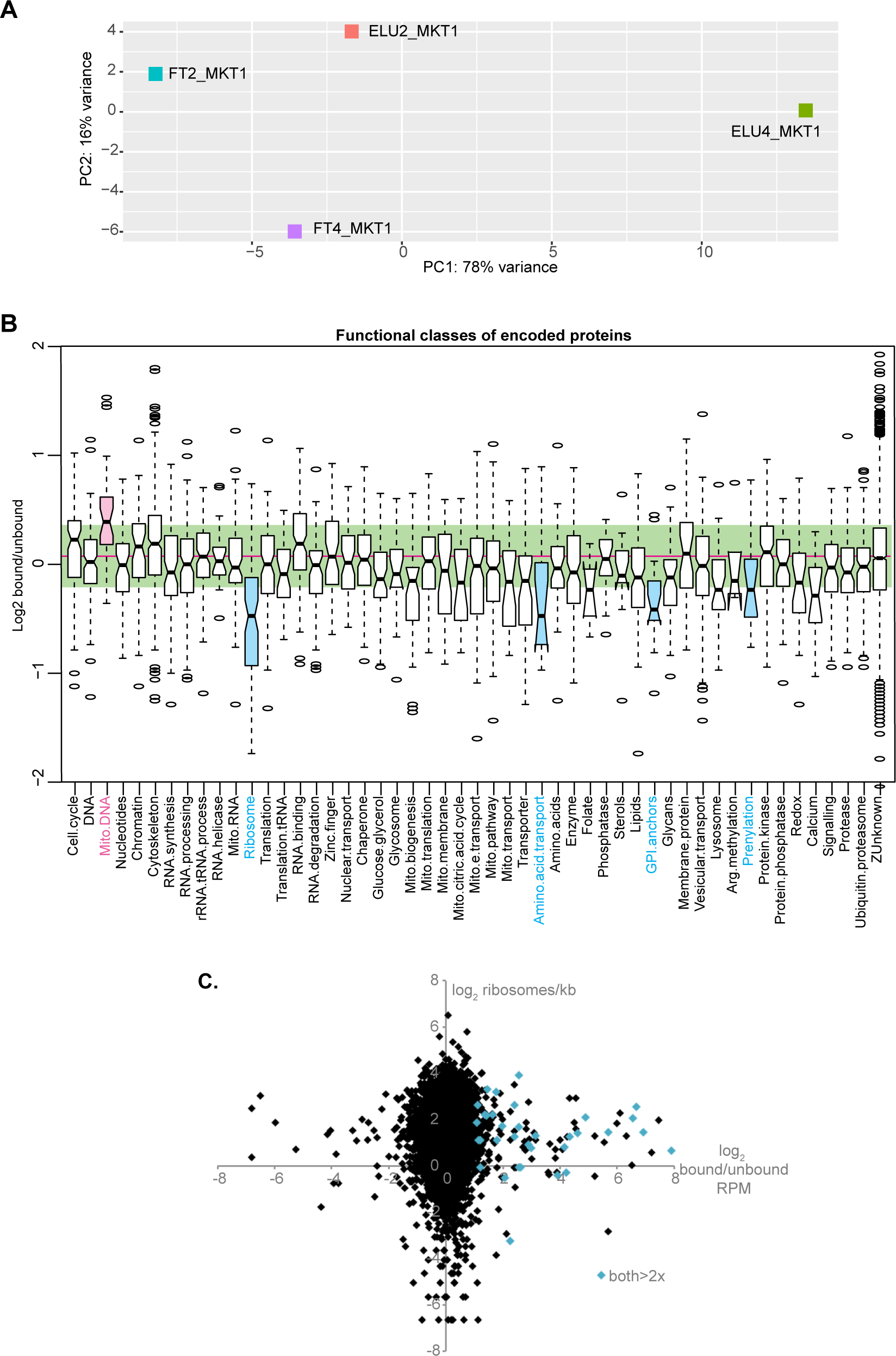
RNA binding by MKT1 and MKT1L. A) Principal component analysis for MKT1 RNA pull-down, made using DeSeqUI (82). MKT1L-PTP was bound to and IgG column, and eluted with TEV protease. The flow-through (FT) or unbound fraction was saved and RNA was prepared. The bound protein was released using TEV protease, then eluted, and RNA was prepared from that fraction as well (eluate, “E” or “bound” fraction). Results for two independent datasets are shown. B) The raw read counts for a set of unique genes were converted to reads per million (RPM) then the ratios of eluate (MKT1-bound) to flow-through (unbound) were calculated. The average was taken then log2 transformed. The Figure shows the distribution of these ratios for mRNAs encoding proteins of different functional classes, as listed in Supplementary Table S3. To make the plot, 206 of 4514 mRNAs were excluded. Of these, 181 were removed because replicates differed more than two-fold, and 25 were excluded because the ratios were outliers (greater than 4x or less than 0.25x) and this made results for the vast majority of mRNAs very difficult to see. For mitochondrial DNA 3 out of 26 mRNAs were significantly enriched in the bound fraction, while 11 out of 71 ribosomal protein mRNAs were significantly depleted. C) Relationship between MKT1 bound/unbound ratio and ribosome density in bloodstream forms (12).

**Figure S9.**
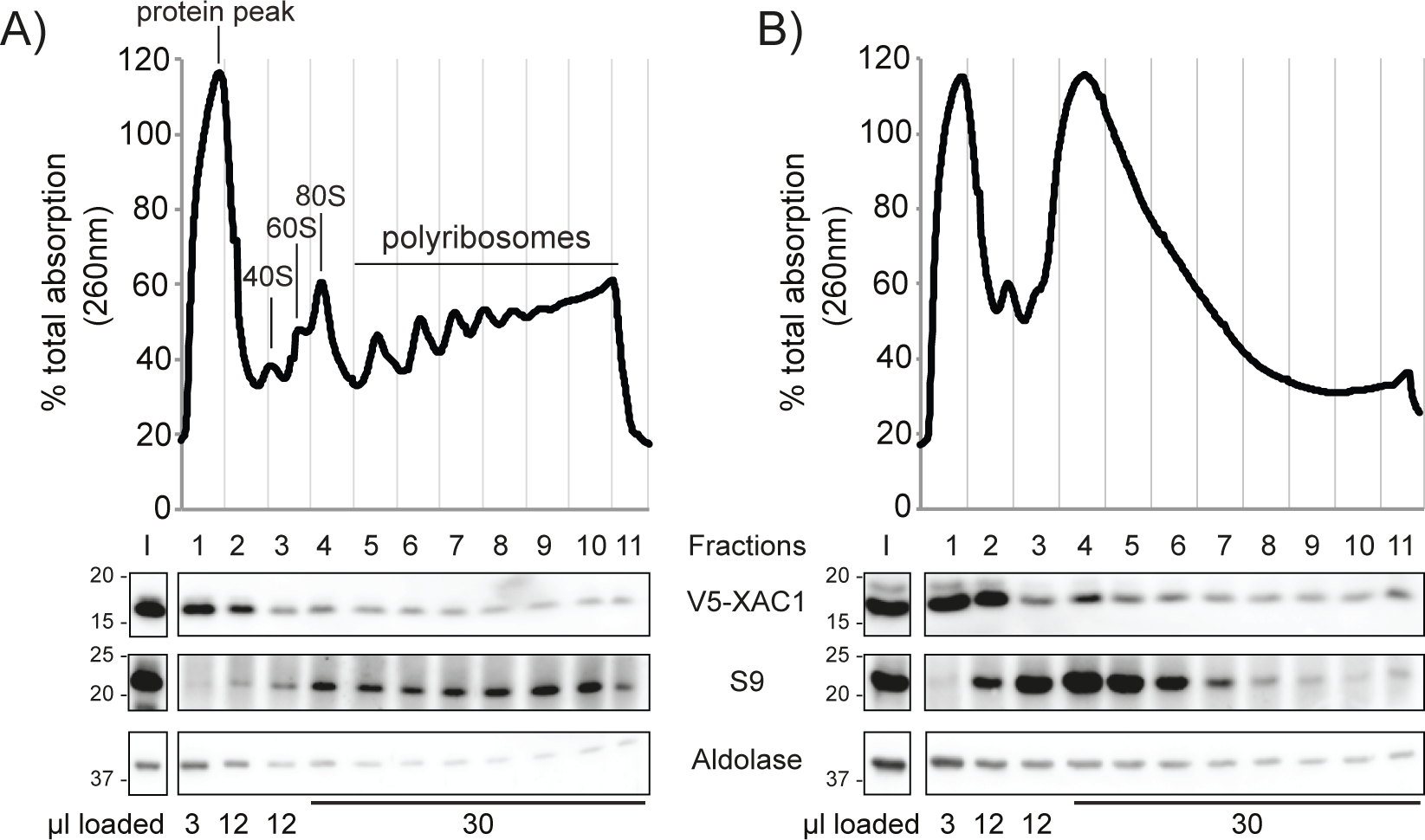
XAC1 is mostly not associated with polysomes. Polysome fractionation was done in sucrose density gradients in the presence of RNAse inhibitors (A) or in the presence of 100 µg/ml of RNAse A (B). The top panel shows a typical optical density profile after sucrose density gradient centrifugation of total extracts from trypanosomes expressing V5-tagged XAC1 briefly treated with cycloheximide. Fraction numbers are bellow. Fraction I is the input and fraction 11 is the heaviest fraction. The bottom panel is the Western blot showing the distribution of V5-XAC1 in comparison with the ribosomal protein S9 and aldolase.

**Figure S10.**
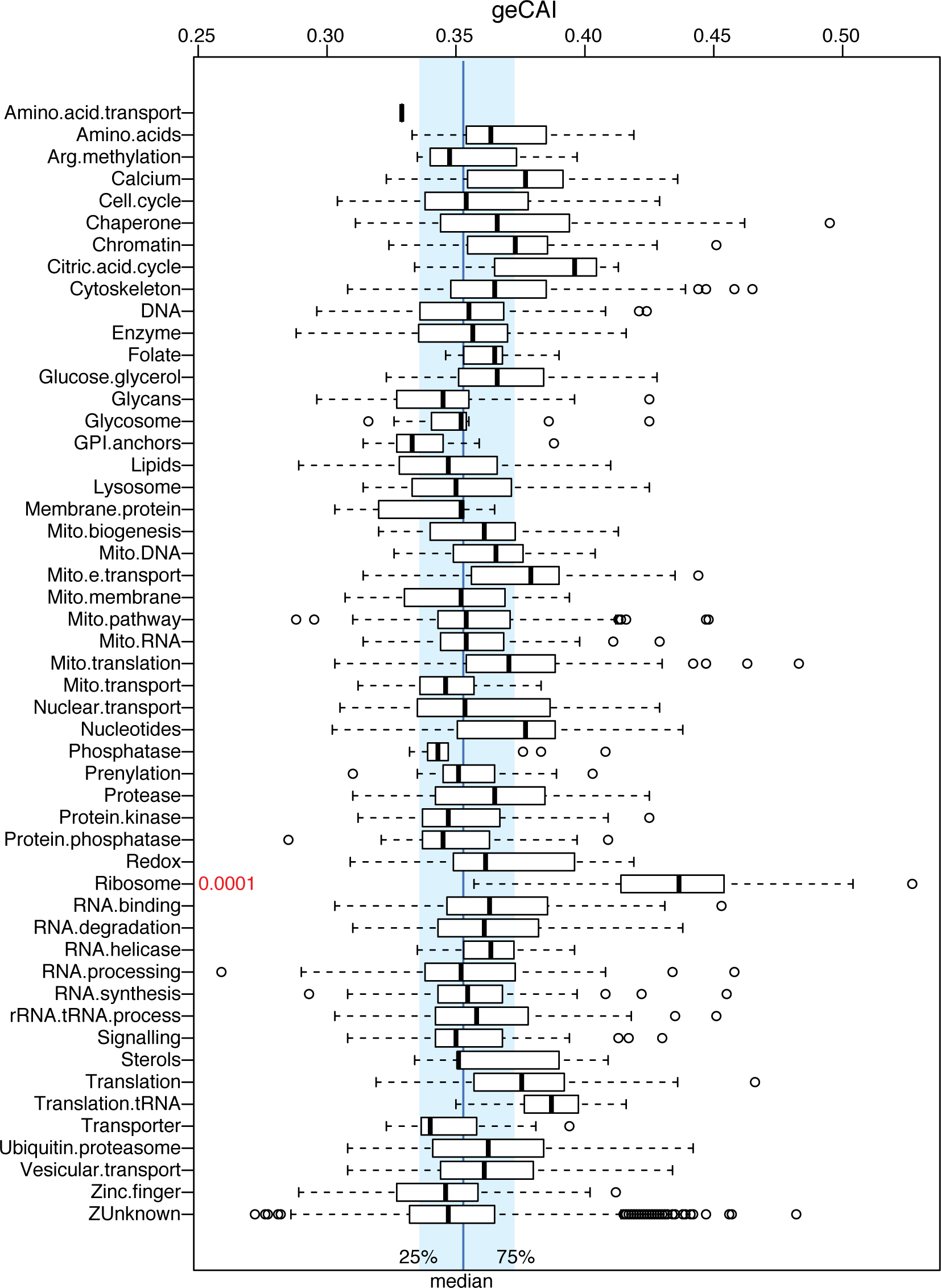
Codon optimalities of gene categories used in this paper. Genes were classified as in Supplementary Table S3, and geCAIs were taken from (8). Results for the functional groups were plotted as in Supplementary Figure S5B. P-values from ANOVA were not corrected for multiple sampling. Since about 50 categories were considered, a threshold P-value for <5% chance is 0.001. The exceptional geCAIs of ribosomal protein mRNAs were already noted by (8); the anlaysis shown here does not reveal any other significant exceptions, although repeated genes (apart from those encoding ribosomal proteins) are missing.

Supplementary Table S1. Plasmids and oligonucleotides

Supplementary Table S2. Proteomic Data.

Supplementary Table S3. RNASeq results

