## Supplementary material for "Essential PBP1-associated proteins of *Trypanosoma brucei*": Figure S3

#### MKT1 & MKT1L alignments



```
TCSYLVIO_007992     ----MRKVGGNRNNER-QGRGQ-GQQGSPMAPPVSSMGYQQHQQQHHQYHHH--------   46
TvY486_1001495      ----MRRLGGNRNSER-QSRGQ-GQQGSPIPAPGPSMGYHQHQQHHHHHHHHHHHHHHHH   54
Tb927.10.1490       ----MRKAGANRNNER-QGRNL-GQQGPQMSPPGPSMMYPHHHQHQHHHQHQHQHQ----   50
TcIL3000_10_1290    ----MKKGGPNRNNDR-QGRNQ-GPQGSQIAPAGPSMVYQHQHHQHHQHQHHHVHHHS--   52
Baya_167_0010       ----MITTD--SEPM-SQLRASPSQVHQQQPRPQETGSS---NQSPPA------------   38
LmjF.21.0820        MPPRMHRGGGGQGPAPQQQQAPPPQQ--Q----QQQAPPQQQQQGYGN------------   42
EMOLV88_210013000   ----MHRAGGGQGSAPQQQQAPPSQQ--PLPQPQQGPPQHQHQQGYGN------------   42

TvY486_1001495      PHQHQHQHQHQHQHQHLQHQHQHQHQHLQHQHQHHQHHHQHHHQHQHQHQHMQHHQHQHQ  114
Tb927.10.1490       ----------------------H-------------------------------------   51
TcIL3000_10_1290    ---------------HQYHQHQHQHQH---------------------------------   64

TvY486_1001495      HQHQHQHQHLQHQHQHQHQHLQHQHQHHQHHHQHHHQHQHQHQHMQHHQHQHQHQHQHQH  174
Tb927.10.1490       ------------------------------Q--------------------HQHP-----   56
TcIL3000_10_1290    --------------------HQHQHQHHQHQ--------------------HQHQ-----   79

PCON_0044260        --------------------------------------------------MPPARRTN--    8
TCSYLVIO_007992     --------------------HHHQHHHNSSYSNV-----------YEGSRPPP-------   68
TvY486_1001495      QHLQHQHQHQHQHQHLQHQHHHYHMHHNSGYNTM-----------YDNMHMHT-------  216
Tb927.10.1490       ------------HQ-HPHQHHHHHPHHNGVYGNA-----------YDNIR----------   82
TcIL3000_10_1290    ------------HH-QHQHQQHQHQHHGGVYGSP-----------YDNMR----------  105
Baya_167_0010       ------------GA-PHPSHFHHFHQHNQPYIANGNNGGYGDHHFNNVYPGPSAQGASG   85
LmjF.21.0820        ------------GTGGPQSHLHVFHHHA-AYNDVGRGGM-----------PPPQHGGNSG   78
EMOLV88_210013000   ------------GAGGPQSHLHVFHHHT-TYNDTGRGGM-----------PPPQHGENSG   78

PCON_0044260        --------------------RGGSGGAAQHQVPFSVGGMAPGGKSYSQAPYSQYGSQPPQ   48
TCSYLVIO_007992     ---------HSMQQMSPHGYGPGGGGGTPAA---AGGN----G--W---------NT---   98
TvY486_1001495      ---------HNVQQVAMHGYGQGV-----------GGN----G--W---------SG---  238
Tb927.10.1490       -----------QPQMPPHSFGQGV-----------SGS----P--W---------NS---  102
TcIL3000_10_1290    -----------QQQMPPHGFGQGV-----------ATS----G--W---------NG---  125
Baya_167_0010       NANANWGNNHSNAYNNSSSPSGGYGSHNLPPPPPQGGG----GGVMPQAQ----------  131
LmjF.21.0820        S-----------NNMSSMMYGGGSGGGNAPTSPVYGGG----GTAYGNNK----SSIPPH  119
EMOLV88_210013000   S-----------NNMNNMMYSGGGGGGNPPTSPIYGGA----GTPYGGNN----GNIPHH  119

PCON_0044260        QQ-PPQQQQQQPQ-------SHH-------------------MQY----GNMLSGQQPPP   77
TCSYLVIO_007992     ---PQQQQQQMP--------MYNQGYGHMGPG--AGGGSG--GGY----GDM--GGVYQD  137
TvY486_1001495      ---PPP--PQVS--------MYNQNYGHMGPGGVAAGGGG--GGY----GDM--GGMYPD  277
Tb927.10.1490       ---PP---QQTP--------MYNQGYNQMSPG------TG--GGY----GDM--GGMYRD  134
TcIL3000_10_1290    ---PP---QQAP--------MYNQGYNQMPPG------GS--GGY----GDM--GGMYPD  157
Baya_167_0010       --------QAYGQG------NMGRGYVTPSPH--QAGGMIQQNQYMNRQGSM-NNRRGAR  174
LmjF.21.0820        MMSPSPHPQIVGMGNEYAPRMQSSPYPQQQ-Q--QGGGASTPNSIGRRVRTPGPSNMGPQ  176
EMOLV88_210013000   MMSPSPHPQMVGMGNEYTPHMQSSPFPQPPPQ--QGGSVSTPNSMTRRVRTPGPSGMGPP  177

PCON_0044260        SMPSQQPPHPQSQNWSS--NAKAPPAPSDYQGNSNPAGGPNVPPQQYGGGGAGYQQPTSP  135
TCSYLVIO_007992     MHPGQYPPSPSPSHMGG-------GSGMPYMRNPPMHGGVPPAGP-IPGPYGGLPGQGAP  189
TvY486_1001495      MHNGQYSSP--QSHMGG-------GAGVPYVRNAPMHGSVPPSGP-HPGGYGGIPGQGGT  327
Tb927.10.1490       VHGGQYPSS--PSHMGG-------GANIPYGRNPAMHNGL-PAGAPLPGGYGALPHQGA-  183
TcIL3000_10_1290    MHGGQYPAS--HSHMGS-------GSGMPYGRNPPMHNGVAPAGPHIPGGYGALPHQGA-  207
Baya_167_0010       SQGANMQPPPQQQAYYPPSSQQTPPPQVNYNNQSGMRGGM---PPHHMEPPLGYPS----  227
LmjF.21.0820        PQQQQLPPPPPQQPYYGGNQGG----GMSY--NPNMRGSAPPPPPPHMEPSPSFVNGTPQ  230
EMOLV88_210013000   PQ-QQLPPPPQQQPYYGGNQGG----GMGY--NPNMRGSAPPPPQPHMEPSPSFVNSGSQ  230

PCON_0044260        LRPP-VS-----------------------------------------------------  141
TCSYLVIO_007992     -------------------------LPP--------------------------------  192
TvY486_1001495      -------------------------PLA--------------------------------  330
Tb927.10.1490       -------------------------HTP--------------------------------  186
TcIL3000_10_1290    -------------------------HSP--------------------------------  210
Baya_167_0010       --PPMMSRADNAFGGHDMGNNSSPPPQG--------------------------------  253
LmjF.21.0820        QPQPGMMRGDGGYGP---DVNNSPPPQNNGYRAGQPPPPPQQHQQ-QMHMQSSPLHPQHY  286
EMOLV88_210013000   QPHPGIMRGDGGYGP---DVDNSSSPQNNGYRGSQPPPPPMQQQMHMHSPSTVPPQQHY  287
```

### MKT1 & MKT1L alignments

```
PCON_0044260          --------GGGVNVSAYNEGYYQSPMPQPISGHGPMYGLQSHLPHHQHQGSRPPSMLPQL   193
TCSYLVIO_007992       ---MGYGRGGHPPMD---------------------------------------------   204
TvY486_1001495        ---MGYGRGGQPPMD---------------------------------------------   342
Tb927.10.1490         ---MGFGRGGQPLME---------------------------------------------   198
TcIL3000_10_1290      ---IGYPRGGQNIMD---------------------------------------------   222
Baya_167_0010         ---GNYGGYGNPNMGDPQGSYYPNPMRAGNNVN---------------GGSPSRSVP---   292
LmjF.21.0820          QGNGGYGGGGGRGMG---------------------------------GGMQNPNMQ---   310
EMOLV88_210013000     QGNGGYGG--GRGMG---------------------------------GGMQNPNMQ---   309
```

```
PCON_0044260          GNGYSMPPPTHQHHQHMQGDYSSQGGHLSGGAPPP-QHHHHHHHAMRGAPAYPVQCQYPM   252
TCSYLVIO_007992       --------------------------------------PMQ-QGGMYGGPMVRHD-GY--   222
TvY486_1001495        --------------------------------------SMQ-QSSMYCGPMMRHE-GY--   360
Tb927.10.1490         --------------------------------------HMQ-PSGMYGAPMMRHD-GY--   216
TcIL3000_10_1290      --------------------------------------SMQ-QGGMYGAPMMRHD-GY--   240
Baya_167_0010         ---QQPPPPPMSSG-----------------NPYNMMEPRNR--NMYNNSMP-PQ-QV--   326
LmjF.21.0820          ---MQP---PPQQQQHMMGGGGG---SASNSANRGMMGPMGGSKQMQQGPPPPLQ-QQ--   358
EMOLV88_210013000     ---IQSPPPPPPQQQHMMGAGVG----APNNSNRGILGPMGGNKHMQQGPPPPPQ-QH--   359
```

```
BODOCUG91050          ------------------------------------------------------------   0
PCON_0044260          ADGMRPSVPGHNMHSSQMVPPMHAH------HLGNQQPP-IQQHQMQQYAEVGVRVAGNQ   305
TCSYLVIO_007992       -------------PDNRISPPP-QVGYGPNVGAPPPPPP-------------------GI   249
TvY486_1001495        -------------PDSRMSPTA-QIAYGPGSAAPP---------------------MPGP   385
Tb927.10.1490         -------------PDTRVSPPA-QVGYGAGAPVGPPPPP---------------PMPAGP   247
TcIL3000_10_1290      -------------PDTRMSPPA-QVGYGPNTPVGPP-PP---------------PMPAGP   270
Baya_167_0010         -------------PPSHQPPAPSQQQP-------------LH----PQQAMQGGPRMGNQ   356
LmjF.21.0820          -------------QQQAMPPPPQQQQPGMNSNWGSPQPPPTHMQANVSPPPQGQPQMGGM   405
EMOLV88_210013000     -------------PQQ--MPPLQQQQPGMNSNWGSPQPPPVHMQPNASPSPQGQPHMGGM   404
```

```
PCON_0044260          PFNGGSGVDVMVRPPHQAGPSALPQDGMGNTYMGRNGAPGFAPGGRTNGG----------   355
TCSYLVIO_007992       PYGG------------------GN-------PMPVARSPAGVP-----TAAGGYPDPRSK   279
TvY486_1001495        PYGGVPAS---------TGPGPGG-------SMPIPRPPG--------GGSGGYVDPSGR   421
Tb927.10.1490         PYGGVLPG---------AGVGPGG-------PMNASRPPAGIGGGGGGGGNAGYLEMRAR   291
TcIL3000_10_1290      PYGGVIAN---------SGVGPVG-------PPGVSRPPAP-------GVAGGYLDMRSR   307
Baya_167_0010         ----------------------------------------------------------DW   358
LmjF.21.0820          QYGGG---------------GPVP-------MVGRGRVPGARMGGGGMGPMGGN----NM   439
EMOLV88_210013000     PYGGG---------------GPVP-------MVGRGRVPGARMGGGGMGPMGNN-----M   437
```

```
PCON_0044260          -AAPVPTPGLGPQ-MQPLPQPQHQHMGQTGYVD-----------SVAHPY----------   392
TCSYLVIO_007992       GMLNNPMPMLNAPRHPRVGTPEPGIMGG-GIPPGRFNHQSALAPP---GMLP--------   327
TvY486_1001495        GMMNSPMSLPSAVRQPRIGTPDAGVV-----PLGRLNQSHPMMPH---GMASS-------   466
Tb927.10.1490         GM-GPQMPASNASRPPRIGTPDPGMM-----PPNRMNQS-PMMGQ---GLPPQ-------   334
TcIL3000_10_1290      GM-NSQMGPSNASRGPRIGTPDPALL-----HHNRMNQS-PMMNH---AMPPQ-------   350
Baya_167_0010         GSPPPM-GQYNNS-RPILPP-QAGHM-------Q----QMNMPPPQVHQMPPQGHPGPA-   403
LmjF.21.0820          GAPPPP--PQQQQRQPPQPPSMMSGMGGP----QQMPGGGNMNPQMMMPPPPQGGQGPMM   493
EMOLV88_210013000     GAPPPPLPQQQQQRQPPQPSSMMGGMGGIGGPQQHMSAVGNMNPSMMMPPPPQGGQGSII   497
```

```
PCON_0044260          -------PTEYRDPMSVPFSAAASPLPADADAVPMMQRAMD-AEASVQRPIADMYHFLVP   444
TCSYLVIO_007992       -------PANYMPVGSVPPNQPPI-----MPAPTSPGDPLHVERFSLGGPTENLYDFLYE   375
TvY486_1001495        -------PNNYMAGMGRSPSSRVI-----ASTPPPVVEPFRGEEDTGRPPTDTLYDFLQE   514
Tb927.10.1490         -------VGNFMPLPGVSPSPHAM-----IPTTPSMGDTFPGDDANMSGSTEALYDFLHD   382
TcIL3000_10_1290      -------VNNFMAMGVQPPSPQVM-----VPTTPSMGDTFPGDEANMGAPIEALYDFLQG   398
Baya_167_0010         --SRGAPPGPM--FGGTYPSPHDV----PANAPPMDADWMP-MMNPAPEMHESFYDFLKN   454
LmjF.21.0820          QQSGGV-PGMY--GGSMD-----ANMMGGMGGPLMPG-YMDADPRGDLAPQQTLYEFLME   544
EMOLV88_210013000     QQGGGVPPGMY--AGGMDPNTISGDMMGGMGAPPMPG-YADVDPRGDLATQQPLYEFLMG   554
TCSYLVIO_003374       -----------------------------------------MYPRQDD--VSLLQRYIND   17
TvY486_0604140        -----------------------------------------MYPRQED--ISLLQKYVNE   17
Tb927.6.4770          -----------------------------------------MYPRHDD--VSLFQRFVND   17
TcIL3000_6_4240       -----------------------------------------MYPRLDD--VSLFQRYIND   17
Baya_226_0010         -----------------------------------MNRVGLVHSRHGGTHPRSFQDFVSE   25
LmjF.30.3430          -----------------------------------------MYPHSKQSMASTIQSFVTE   19
EMOLV88_300040300     -----------------------------------------MYPHTKQTMALTIHSFIAE   19
```

PIN domain

### MKT1 & MKT1L alignments

```
                                          1   2
PCON_0044260          RDLVERTQLSNFFPENYDAARDGPLKLAMDGNYCLGWLKDKLHAHDPLWFLHSSLPDELL   504
TCSYLVIO_007992       HDLVSVDNLSSFFPEGYDPKRDPPLKIAVDGNFCLNSLRDELRKRDPLWFLHSTLPEELL   435
TvY486_1001495        RGLVSIDNLSRFFPERYDMKNDPPLRIAVDGNFCLTSLRDELKKKDSLWFLHSTLPEELL   574
Tb927.10.1490         RGLVSVDNISKFFPEGYG-KDDPALKVAVDGNFCLTSLRDELRKRDSLWFLHSTLPEELL   441
TcIL3000_10_1290      RGLVSVDKLSYFFPEGYK-EGDPRLKVAVDGNFCLTSLRDELRKRDSLWFLHSSLPEELL   457
Baya_167_0010         RMLLKTCPLSDFLPTCSDPSATGAVTIAIDGNYMISSLQDRLQEIDLLWFLHSPLPDVLL   514
LmjF.21.0820          KRLLRKTTLGTFFPEGYDPKCNSRITIAVDGNYMITNLRAQLQRTDPLWFLHSCLPDCLL   604
EMOLV88_210013000     KRLLRKTTLNTFFPEGYDAKRDGRITIAVDGNYMITNLRAQLQRTDPLWFLHSCLPDYLL   614
TCSYLVIO_003374       NKLEEKGV---PLTELRRLEGTGQVMLGVDGTKIIDLITQAVREKEKMAIYTYTTPFTVY   74
TvY486_0604140        NRLEEKGI---PLHELSRLEGTDQIMLGVDGTKVIELITQAVRDREKMAIYTYTMPFTVY   74
Tb927.6.4770          HKLEEKGL---PLYQLAQLEGTDQVMLGVDGTKIIDMITQAVREREKMAIYTYTTPYTVY   74
TcIL3000_6_4240       NKLEEKGI---PLHELTPLEGLNKVMLGVDGTKIVDTITQAVREKERMAIYTYTTPYTVY   74
Baya_226_0010         NNLEEKGI---PLNKLARLPETNQVVLAYDGNKILDVITEAVREEEKMSIFTSSTPLTVY   82
LmjF.30.3430          TKIEEKGL---KLSELRRLPNSNQVSIVFDGNRVLDDVVQDVHVTGPLTIFTASTPWSAY   76
EMOLV88_300040300     NRLEERGL---KLSELRRLPNSNQVSIVFDGNRVLDDVVQDVHATAPLTIFTASTPWSAY   76
```

```
                                                3
PCON_0044260          DLLRENM-ESLRS---------VNLEPILVFNGLGISGDVEAYLMTQDELTSRSTV----   550
TCSYLVIO_007992       DSVCHHV-EWMRS---------LNLEPIWVFNGLSVSGDVETFLTSEGELRARDIV----   481
TvY486_1001495        RLVWHHV-EWMRS---------LKLEPIWVFNGLSVSGDVETFLTTEAELRARDRV----   620
Tb927.10.1490         MLVQQHV-EWMRN---------MKLEPIWVFNGLSVSGDVETFLTTEAELRARDAV----   487
TcIL3000_10_1290      VLVWHHV-EWMRN---------LNLEPIWVFNGLSVSGDVETFLTTENELRARDAV----   503
Baya_167_0010         SLIEENV-KRMRE---------KNLEPIWVFNGVGIGGDVEAFIPSTKEGIARNLA----   560
LmjF.21.0820          SLVHKHV-EQMRA---------HHLEPLWVLSGISINGDVEAFLPSPDELHSRDAV----   650
EMOLV88_210013000     TLVHRHV-EEMRS---------NNLEPLWVLNGISINGDVEAFLPSTDELHSRDAV----   660
TCSYLVIO_003374       DKITDVC-KTLRSL--------KNCTPVLVFNGIPFYPDPSDDNCREKNMVPPDVAALNG   125
TvY486_0604140        EKIDEIC-RMLRSI--------RNCTPVFVFNGIQFCPDATEEFSKEKIAAPIEVATLIG   125
Tb927.6.4770          EKINEFR-TMFQSI--------KNCTPVFVFNGIQYSPDSAEEFGREKNVVPSEVAALTG   125
TcIL3000_6_4240       EKVDEIR-TMLRSL--------KNCTPVIVFNGIQYSPDPAEELSREKNMVPVEVAKLVG   125
Baya_226_0010         RLIKRYC-EQAEAIGRHFESHDAVFKPVFLFNGCGFMPDADLDLTL--PTQPNDVAHCRA   139
LmjF.30.3430          TIISEYTzEMVRKLGEYFRACDADFKPMFVFNGCGFHPMEDQDTTS--PAPPMEVASYSA   134
EMOLV88_300040300     TVISEYT-EMVSKLEEYFRAFDADFKPMFVFNGCGFHPMEDQDTTS--PTPPMEVATFNA   133
```

```
                                                                           4
PCON_0044260          --WKTLLNGE-L-PDDELS-RSAFDQ----SLGEDVHMAILRFLQKELGVMVVAAPFLNW   601
TCSYLVIO_007992       --WSRLEEGE-I-PDEADI-QTAFDQ----PLGEDVQMAVSRYLKKELNVMAVTAPFLNW   532
TvY486_1001495        --WCELEDGL-I-PSETEI-QEAFDQ----PLGEDVQMAVARYLRHKLNVMVVTAPFLNW   671
Tb927.10.1490         --WSKLEDGE-I-PDEVEI-QEAFDQ----PLGEDVQMAVARYLKEELGVMAVTAPFLNW   538
TcIL3000_10_1290      --WSKLEDGS-L-PEDMEI-QEAFDQ----PLGEDVQMAVARYLKDKLQVMAVTAPFLNW   554
Baya_167_0010         --WDRLMEGFLL-T-PGEA-AEAFETS--SSIGEDVLMVVQRFLRSQLGVLTVTAPFLNW   613
LmjF.21.0820          --WAKLNEGIDL-PTQDEI-TEAFETS--SAVGEDVLRAIQRFLRTEENVVCVTAPFLNW   704
EMOLV88_210013000     --WSKLNEGIEL-PTQDEI-TEAFETS--SAVGEDVVRAVQRYLRTEENVVCVTAPFLNW   714
TCSYLVIO_003374       TDSSRLCN-MHS-MRAVDI-QKKSSSR--FFVEEDVENQIVKLFRSEFK-DTMRAPYLAW   179
TvY486_0604140        TDSSRIAN-MPN-GRQMDI-QKKVSNR--FVVEEDVESQIIRLLRTWFK-LIMRAPYLAW   179
Tb927.6.4770          TDSSRLSN-TTN-VRFAEI-HKKTANR--FVVEEDVEGQIIRIFRSEFK-NTIRAPYLAW   179
TcIL3000_6_4240       TDSSRVD--KSN-IRFLDI-QKKAAAR--FVVEEDVEGQIIRILRHEFK-NTIRAPYLAW   178
Baya_226_0010         FLLQ---T-TVK-NTPLEI-QRQFAAR--FSIDDDVEGLIVRCFHAKFT-DTFCAPYHSW   190
LmjF.30.3430          YNKN---K-TVR-SVPAEA-QKKFASR--FAIEEDVEGLIVRKLCEVAE-ETMRAPYLAW   185
EMOLV88_300040300     YSKN---K-LVR-NVPIEA-QRKFACR--FVIEEDVEGLIVRKLCEVAE-ETMRAPYLAW   184
```

```
                      5          6
PCON_0044260          AQMVTFVEEGV--ADLLFGPSEALCIPYDPMKLIIDIELTNGTPVVAFLNRDRVLRTLFP   659
TCSYLVIO_007992       AQMVAFHKEGL--ADLLMGPPEVLLLPYDNMKLIVQIDVSNS--NVSYFDRERVLRVLFP   588
TvY486_1001495        AQMVAFHKEEI--ADLLMGPPEMLLLPYDEMVVIVQIDVSNN--NVSYLDRDKVLRALFP   727
Tb927.10.1490         AQMVAFHKEGI--ADLLMGPPEMLLLPYDEMKVIVQIDVSNS--SVNYLDRDRVLRALFP   594
TcIL3000_10_1290      AQLVAFHKEGI--ADLLMGPPEMLLLPYSDMKVIVQIDVSNN--NVSYLDRDRILRALFP   610
Baya_167_0010         CQMATFHVEGD--AALLMGPPEMLLVPYEGMKLIVEMDLGSD--TVAYFERDHVLRQLFP   669
LmjF.21.0820          CQMCTLHKEGI--AQLLMGPPEMLMVPYDDMKVIAEVNLQTS--EVAYYDRDEVLRELFP   760
EMOLV88_210013000     CQMCTLHKEGL--AHLLMGPPEMLMVPYDDMKAIAEVNLQSG--EVAYYDRDEVLRELFP   770
TCSYLVIO_003374       AQLSAFCHPKNRHISEVYGCLELLAFPGID-RVITNINVMKG--TVDMVRKSRLLELLR-   235
TvY486_0604140        AQLSAFHHPSNRHVSEVYGCLELLAFPGIE-RVITNINTANG--TFDMVRKSRLLETLH-   235
Tb927.6.4770          AQLSSFRHCNHRHISEVYGCLELLAFPGID-RVVTNINPTRG--TFDVVYKARVLEAAR-   235
TcIL3000_6_4240       AQLSSFRHGSNRHVSEVYGCLELLAFPGID-RVVTNINTANG--TFDVVHKSRVLGSLR-   234
Baya_226_0010         SQISAFFAPSNRYVSEVYGCLELLAFPGID-RVVTNINRENG--TFNSVRKSSVLAALRK   247
LmjF.30.3430          SQISAFFAPSNTMTSETFGSLELLAFPGID-RVITNIDVDAG--TFDCVSKASMMAALRR   242
EMOLV88_300040300     SQISAFFAPSNTMASEVYGSIELLAFPGVD-RVITKIDVVAG--TFDCVSKAAVMSTLRR   241
```

**(1)** D->G in *S. cerevisiae* Mkt1 affects gene expression, gives better growth at 41°C and better sporulation. **(2)** S288C *S. cerevisiae* has D->A. This inactivates Mkt1. **(3)** This is D in XPD1 and Fen1 endonucleases. D->A is inactive, D->E is active. **(4)** This is E in XPD endonuclease. Mutation to A: can still bind DNA but is inactive. **(5)** A->V in XPD has reduced activity. **(6)** In XPD, D->A abolishes 3' excision.

MKT-N domain

### MKT1 & MKT1L alignments



```
PCON_0044260        HHVTETDTRAASDRFIDFCLITASHPALSSARVALRMSIQDIYEELSTTAPHYPSICKML   719
TCSYLVIO_007992     YHVTEISTRGAGDRLMDLGLITATHPALSSTSVGLNLTMEEVYEELSSETPKFRSIKDFI   648
TvY486_1001495      HHVTESSTKLAGDRLMDLGLIM--PCALSSARVNLELSMQEIYEELSALTPKYPSIKEFI   785
Tb927.10.1490       NHVTETNTRVAGDRLMDLGLITATHAALSSARVTLNLSMQEVYEELSTPTPKFRSIKDFI   654
TcIL3000_10_1290    HHVTETSTRDAGDRLMDLGLMTAMHPALSSARVNLDLSMQEIYEELSSKTPKYRSIKDFI   670
Baya_167_0010       KWVRGDSTEVAGDRFIHFGLLITSHPAITAARVTIPLSATEIYEELAATNSRTQALQRFI   729
LmjF.21.0820        RYVTETSTAAAGDRFLDFGLLIASHPAITTAHASVQLSTQSIYEELSTPHPNFNTLREFI   820
EMOLV88_210013000   RYVTETSTAAAGDRFLDFGLLIASHPAITTAHASVQLSTQSIYEELSTPHPSFNTLREFI   830
TCSYLVIO_003374     -----ISEDD-------LGSLIVVDSR------------NRVMRTVGPKFSSFDDMCKKI   271
TvY486_0604140      -----VSEED-------LESLIAIDSR------------SRPMKPVALKFQNFEDMCKKI   271
Tb927.6.4770        -----LSEED-------LSSLILVESR------------SRVMRTVTLKFSSLEDMIKKV   271
TcIL3000_6_4240     -----LSEED-------LASLILLDSR------------SRVMKPVALKFSSLEDMCEKI   270
Baya_226_0010       KVG-DITDEE-------LSAFILMDSR------------NKQFRVP-GHLVGFDELCTTG   286
LmjF.30.3430        RYDAEFSEDD-------FSAIALYETK------------NRLFRVK-NRRETFDDLCRLP   282
EMOLV88_300040300   RYDAEFSEDD-------FSAITLYDTK------------SRLFRVK-NRRESFDDLCRLP   281
```

```
PCON_0044260        GDQSSEASEPSEHGQRC--RTAVCLKHAKGRNYIRYNVVFSTKQ-KDCPLTYLR-MVLEP   775
TCSYLVIO_007992     NAHAYVQ----DANKK---KAGLIIKHSKGRGYLRYSGVFSSKL-QDSPLVYLI-RVLEP   699
TvY486_1001495      GKHSRGQ----DSN-K---RGGLAIKHSKGRGYLRYSPVFSSKY-TESPLVYLV-RVLDP   835
Tb927.10.1490       NAHACSQ----ETG-K---KAGLSIKHSKGRGYLRYSAVFSTKS-RDTPLVYLV-RVLDP   704
TcIL3000_10_1290    NAHASTQ----ETN-K---KAGLTIKHSKGRGYLRYSAVFSSKT-PESPLVYLV-RVLDP   720
Baya_167_0010       FLREMCF----APPERR--PGDELLRLARGRCYLQYSAVFSKQT-PECPLVYFK-RVLDP   781
LmjF.21.0820        DQYEYPQ-----NQNER--PNLEKLKHSKGRTYIQYSPVFSNQVTSESSLVYFK-RILDP   872
EMOLV88_210013000   DQYEYPQ-----SQNER--PNLEKLKHSKGRTYIQYSPVFSSQVSFDSSLVYFK-RILDP   882
TCSYLVIO_003374     TRVNDLSLGASYVHQLHQEAMHL-SEQARNRTLNLKSAVLRNLAALGCPVLTLVPPYCTL   330
TvY486_0604140      ERYKDFSLGVSYAYYLHDEVGRA-PETNRSRLQSQRNAAFRNLAALSSPVLTLTAPYCVP   330
Tb927.6.4770        VRYKGSTIGASFAQQLHEEAIRA-SEANRMRASNQRGAAFRNLSAFASPVLTLTPPHCLP   330
TcIL3000_6_4240     VRYKGSTLGASYAQHLHDEVSRGVADMNRAPINNQKSAAFRNLSALASPVLTLTPPFCIP   330
Baya_226_0010       R-WENLQRHTSYLQN-------------SRISEARRKLLRRNIAALDSPVLLKD-GKCIF   331
LmjF.30.3430        R-D--PTRVDAFMKQ-------------HRVGEDQKLLLRRNLGALDAPVFTVD-AQLVP   325
EMOLV88_300040300   R-D--PTRVDAFMKQ-------------HRFSDDQKVLLRRNLGALDAPVFTVD-AQLVP   324
```

```
PCON_0044260        HITLS----DIPVNLVGVFGSHVPVSLFFFHYSGLLSLSIMTVLTQTYLRDDCPVADTDE   831
TCSYLVIO_007992     GLTNA----DMPTNLIGVLGNLVPLSLFYLQFVGLLSVRIMTVITQSYIRDECPVSDTKD   755
TvY486_1001495      NLTNA----DMPANLVGVLGHNVPLSLFYLQFAGLLSVRIMTVITQSYLRDECPVSDTKD   891
Tb927.10.1490       DLTNA----DMPTNLAGVLGHLVPLSLFYMQFSGLLSVRIMTAITQSYLRDECPVSDTKD   760
TcIL3000_10_1290    TLTNA----EMPTNLAGVLGHFVPLSLFYLQFSGLLSVRVMTAITQSYLRDECPVSDTKD   776
Baya_167_0010       TLTNA----NMPNNLSGVLGNLVPLSLFYFQFSGLLSVSLMTIITQSYFRDELPLSDTID   837
LmjF.21.0820        SLTNA----NMPNNLSGVFGYLVPLSLFYFQFTGLLSVGLMTAVTQLYIRDEFPVADTEE   928
EMOLV88_210013000   SLTNA----NMPNNLSGVFGYLVPLSLFYFQFTGLLSVGLMTAVTQLYIRDEFPVADTEE   938
TCSYLVIO_003374     LTRLYESRRQLPIDPKMVMGCPLPPVMYYMFTAGLLSPSLFGALCQGSLVDDWPLVDSIK   390
TvY486_0604140      LHSLYKPHRPISPDLQSSLGMPLPSILYYLMSAGLLSPSLFGALCQESVVDDWPLVDSIK   390
Tb927.6.4770        LHFLYGIPGFPHADAESFVGLPLPPVLYYIMSAGLLSPSLFAAVSQETVVDDWPLVDSIK   390
TcIL3000_6_4240     LHALYEIHGLIPLDVQVFLGFPLPSVLYYLISAGLLSPSLFAAVSQEAVVDDWPLVDSIK   390
Baya_226_0010       LSSVYNRPK--NSDVGTAFGRPIPSIFYYFFMGGPVMPAPIAAVAHSALADDWPLVDTSD   389
LmjF.30.3430        LSTLHNRPR--RCEIRPIFGSPMPIVFFYLFMSGPLLPLPFAIHSQEVLADDWPLIDTSA   383
EMOLV88_300040300   LSTLHNRPR--RGEIRSIFGSPMPIVFFYLFMSGPLLPLPFSIHSQEVLADDWPLIDTSA   382
```

```
PCON_0044260        YHNILHRLMALRAQIVHQLFPRIHNEEYQKRLHELSWVRWYEPILAPF-----------Q   880
TCSYLVIO_007992     YHEKLEVLMTMRSQIIAQVFKKISKDRHPNNTECLSWVRWFKPILAPM-----------Q   804
TvY486_1001495      YHDKLDVLMTMRSQIIAQISRKIGQQSSLKQNECLSWVRWFQPILAPM-----------D   940
Tb927.10.1490       YHTTLGLLMTMRSQVIGQILKRIAHPPPIKRTECLSWVRWFQPILAPM-----------D   809
TcIL3000_10_1290    YHD---------------------------------------------------------   779
Baya_167_0010       YRRYIPTLVALRGQIVFQVVRLMRSETYRRRFERISWVRWYKPLLIVV-----------S   886
LmjF.21.0820        YHHLLHPLMALRGQIISQMVSRIKADAHGQRLGKISWVRWFDSILMSV-----------L   977
EMOLV88_210013000   YHHLLHPLMALRGQIISQMVSRIKPDTHGQRLGKISWVRWFDSILMSV-----------L   987
TCSYLVIO_003374     YRDVAETVLPLRVQTLHQLAASLRM-----SDFGMTWFRRYNAF---------LSRVSKV   436
TvY486_0604140      YRDVAETVLPLRVQTIYQLAWSLRR-----SPGKMSWYRRYNVV---------STRVSKI   436
Tb927.6.4770        YRDVAETVLPLRVQTIYQLAWTMSR-----GLGSISWFRRYNVL---------PARVSKL   436
TcIL3000_6_4240     YRDVAETVLPLRVQTIYQLAWVLRR-----SLGSISWFRRYNVR---------QSRVNK-   435
Baya_226_0010       FRRVAEVILPLRVQIIFQLMQ-FAR-----VDCRMRWLRQYVVFRGT---PNDDHRWSSV   440
LmjF.30.3430        YRRAAESILPLRVQVVFQLFP-IAP-----KTSDFYWIRQYVVFKKQQQAPESQSRVCKL   437
EMOLV88_300040300   YRRAAESILPLRVQVVFQLFP-IAP-----KTSDFFWIRQYVVFKKQQQAPDSQSRICKL   436
```

MKT-C domain

```
PCON_0044260          RLKSPIELDEWDLHGSEAVMQISMDDMTSQGILHTLEISTKVNCFPLGHS--SNTGEGVE   938
TCSYLVIO_007992       RPRDLINLDEWDISSSEEVLQLDEEHLERYSIASVLTFSAEAAR--PIRDQNICSPRETP   862
TvY486_1001495        RPRDLINLDEWELKDNEEIKKLDENRLDDYSIASVLSFTTEAAR--PVNVDGSISPRNMP   998
Tb927.10.1490         RPRDLIDLDEWEISDSDQLKKLDEDCLADYSIASVLSVTADASR--PAVEESNRPAGRVP   867
Baya_167_0010         RPPETILLDEWQLDGCAELVKICEEPLETAPIFTVLSMNSTEICISAAKAI-EANDGKRP   945
LmjF.21.0820          PPDRLIVLDEWNLKNSPELASVPDDQLENIDMSMVLSLKSSVMCIPAPQLPANASPRDAP   1037
EMOLV88_210013000     PPDRLIVLDEWNLEDSGELAGVPDDHLEKIDMAMVLSPKSAAMCIPAPQLPPNASPRDAP   1047
TCSYLVIO_003374       HAPPEIGLDSWNLAGET--------ISENLFLVDVMEFSH-------------LAVSSRH   475
TvY486_0604140        HAPPDIHLDGWNLHNIN--------IPLGLHLVDVMEYAH-------------LACPLEH   475
Tb927.6.4770          HVPPAIQLDGWALHSVT--------VPRGLHLVDVMEFAH-------------LACSVDQ   475
Baya_226_0010         SEPPDISLSAWEVYRPRYM-SSN--EVDNIYFLGVLRFAE-------------CAV-TGD   483
LmjF.30.3430          SNPPTIDLAYWSFMEEELLQSTR--DADNVYFTDIVSFCG-------------CAV-AEP   481
EMOLV88_300040300     SNPPTIDLAYWSFPEEHLLQATR--EAENVYFTDIVSFCG-------------CAV-AES   480

PCON_0044260          KRYNGI-TETYLAILLKVFDFLGYFSHLIKEDQGSQDPHEVE--VL------SDNVN---   986
TCSYLVIO_007992       IRYHGK-RETFFTVLLKAFDFLGYFSHSTVSPEMGEDGEMMA-------GGVEEGVEAVG   914
TvY486_1001495        VMYHGK-KETLYAILLKVFDFLGYFSHSTSPQDTLDEAGAEVEEGVLEYTGHEDNGDGQG   1057
Tb927.10.1490         IRYNSK-RETFLAILLKSFDFLGYFSHSTAPNDAVDGMEMECC----GMDGHDR------   916
Baya_167_0010         ILYHGQ-KQTFLAVLLKAFDLLGYFSHTVIQDC-DDVPIEDP-DAI---------LDGSG   993
LmjF.21.0820          ILYHGK-KETFFAILLKTFDFVGYFSHSADPMDGAELASEPV-QVM---------VQG--   1084
EMOLV88_210013000     ILYRGK-KETFFAILLKTFDFVGYFSHTADPMDAADLTAESA-QVV---------GQG--   1094
TCSYLVIO_003374       IIYETA-EETCAAVLLQSLDLLGYLTHETRDDGEDVQVSEPS------------------   516
TvY486_0604140        VVYQTV-EETYTAILLQSLDLLGYLTHETQEQLGGGQSSEPS------------------   516
Tb927.6.4770          VIYNTM-EETYAAILLQSLDLLGYLTHETQDLHEEGQSSEPS------------------   516
Baya_226_0010         VLYRDV-NTTVSAVLLRSLDLLGYFTHSTD----AMDCSGSS------------------   520
LmjF.30.3430          VIYKSV-QATLAAVYLRSLDFLGYFTHATD----GAESSGPS------------------   518
EMOLV88_300040300     VIYKSI-QATLAAVYLRSLDFLGYFTHATD----GAESSGPS------------------   517

PCON_0044260          -----LAV---------------SANHDYMPQGDESGQVMVPDAGESFICFSKFLTTALR   1026
TCSYLVIO_007992       QGNN-------AEGAPILISDSNSNNYNE----LHDSLPEEGINEYPTVYFPTYLMNAIK   963
TvY486_1001495        QGRSSMTVGGGGGLAPPACFPGGAA-VNEVGDRNSRV-MASEYNGGSNVYFPTYLMATIK   1115
Tb927.10.1490         -----------GGSVPAAYHEKDLSSMKRSEASDINFMADEGLKDYPSVYFPIYLRATIK   965
Baya_167_0010         EHNY-----------------DN-SNDN--------EGTEPGDTPGTTNFFTVYLCRSLE   1027
LmjF.21.0820          QSAP-----------------DDQSNGPDGGKGDLADSSSDPNMSEQRVFFTAYLSISLE   1127
EMOLV88_210013000     QPAN-----------------EGPPNVSDSASGHLTDSTADPCAGEQRVFFTAYLSISLE   1137
TCSYLVIO_003374       ----------------------------------------------------PF-GRALK   523
TvY486_0604140        ----------------------------------------------------AY-GRALQ   523
Tb927.6.4770          ----------------------------------------------------TF-GRALQ   523
Baya_226_0010         ----------------------------------------------------VY-TRALE   527
LmjF.30.3430          ----------------------------------------------------VY-CKALE   525
EMOLV88_300040300     ----------------------------------------------------VY-CKALE   524

PCON_0044260          -ANPPELHGALVLLTELLRVSILNHRMYTYAQNSPGF--VEDDLGEEDEDPAVILASRIA   1083
TCSYLVIO_007992       -HNPRSFQNAFVVLTEILRVGIIGSLPCRYIDPADPQLAISMENQESNCDPRVLLASRIG   1022
TvY486_1001495        -ANPSSFQGPFVILTELIRVGIISSMPCRYINPANQQEAVPLEDQDDNCDARVLLASRIA   1174
Tb927.10.1490         -ANPLDVQASFVLLTELVRVRIINSNPCRYINPANQQVEISMDDQDTNSDSRVLLASRIA   1024
Baya_167_0010         -QCRESFHSALVRFTELVRVGIINASRYSYEEPKKGM---MPAPNDSDEPAEVLLASRIA   1083
LmjF.21.0820          -HCPQEFQCALVRFTELVRVSTINSIAFTFADTIELQ---HDEDGALADPPEVLLATRIA   1183
EMOLV88_210013000     -HCPQEFQCALVRFTELVRVSTINSVAFTCADSIELN---SDEDGALADPPEVLLATRIA   1193
TCSYLVIO_003374       LCGVPTLSEYVVLLIELSRTGAITTEQFRMT---------SEEIIPRDIPPEIVLASRIL   574
TvY486_0604140        LCCVPTLSEYTVLLIELVRTDAISAEPFRIT---------TDEVHPRDIPEDIVLASRIL   574
Tb927.6.4770          LCSVPTLSEYTVLLIELARTNAISTEPFRIT---------TEEVSPRDTPRDIVFASRVL   574
Baya_226_0010         KFTCPTLSEYGLLLIELIRTGTLTDTPLEVS---------MAHAS-ERYPPGLRFASRLL   577
LmjF.30.3430          RFDCPTLSEYGVLLIELMRTGTLNDDPLVVV---------LNRVD-DTIPMGVRFASRLV   575
EMOLV88_300040300     RFDCPTLSEYGVLMIELMRTGTLNDDPLVVV---------LNRVD-DSIPMGVRFASRLV   574

PCON_0044260          CLLPVKYKSQN-PGEEFSWAPVYSRHLCAFVVMVRAMNRGLRDLVEVITASVHLDGTSDC   1142
TCSYLVIO_007992       CLINLPYRRKT-EDLPFIWAPVYSRHLCAFTVMVRAMCRCLRELVEVIATTVFLAGNSSC   1081
TvY486_1001495        CLVTLPYHRSS-EDLPFV*APVYSRHLCAFTVMVRAMCRSLRELVEVIATTVFLTGNSDC   1232
Tb927.10.1490         CLVKLPYRRAS-ENLPFVWAPVYSRHLCAFTVMVRAMCRCLRELVEVITSTVFLSGNSSC   1083
Baya_167_0010         CLIATPYIDPDMLEQDFQWAPVYSRHLCAFSNMARYMNRYLRELVEVISATVFLNEFSDC   1143
LmjF.21.0820          CLVSVPYQQKTDEESEFEWAPVYSRQLCAFTVMSRVMNRSLRVLTEAIAAALFLSGFSDC   1243
EMOLV88_210013000     CLVSVPYQQQVDEKSQFEWAPVYSRQLCAFTVMSRVMNRSLRVLTEAIAAALFLSGFSDC   1253
TCSYLVIO_003374       SIIPLNV--------SSSWSGPIDPELAAFSMISRMISRSIRHLLEAMLAIIFYQGRTLV   626
TvY486_0604140        SIIPLNV--------SGPWTAPIDAELAAFSMLSRMVSRSIRQLLEAITALMFSKGRTTV   626
Tb927.6.4770          SIIPLNV--------SGPWTAPIDAELAAFSMLSRMISRSIRQLLEAITTLMFSKGRTHV   626
Baya_226_0010         SIIPLNL--------NSAWMGPFDAEMAAFGMASRMLSKTLRYLTETVACLLFATKSTKV   629
LmjF.30.3430          SIIPANV--------CGPWSGPFDPEMAAFGVISRLFSRTLRVLHEVVAMLLFANRATTV   627
EMOLV88_300040300     SIIPANV--------SGPWSGPFDPEMAAFGVISRLFSRTLRVLHEVVAMLLFANRTTAV   626
```

```
PCON_0044260         TLQEFAALSLLLPFGTVPTAIGGLLLHYVLVFPLDYEDYRHTPEGRIAFLQEKFRNIADV   1202
TCSYLVIO_007992      SLVDFSSFAFLLPFGEIPSAIGGLLLHYVLVFPSDYQVNLTTRQERLEYLQEKFRDIPDL   1141
TvY486_1001495       GLEDFAKFSAILPFGDVPSTVGGLLLHYVLVFPSDYQNGLTTAQQRIEYLQGKFRDVPDL   1292
Tb927.10.1490        SLQDFAEFASILPFGDVPSTIGGLLLHYVLVFPSDYQANLTSREERIEYLQGKFRDIPDL   1143
Baya_167_0010        GLDDFASFSPYLPFGDVPSSIGGLLLHYVLVFPADYEHNRNTPKERCDYLQGKFRNIPDL   1203
LmjF.21.0820         SLEDYDSMTPLLPFGDVPSSLGGLLLHYVLVFPPDYEANCQTPEERCRLLETKFKAIPNI   1303
EMOLV88_210013000    SLEDYNSMTPLLPFGDVPSSLGGLLLHYVLVFPPDYEANCQTPEERCHLLETKFKAIPDI   1313
TCSYLVIO_003374      PLHRICAIQQALPFSTPVEFGGGVLIEYVLMKEKCTLSD----------IEAAFPECNYL   676
TvY486_0604140       PLQKVCDIQSMLPFSIPVEFGSGVLIEYMLMKDRCTMSD----------LEMAFPECTYL   676
Tb927.6.4770         PLHRIGEIQRLLPFSTPVEFGCGVLVEYMLMKDKCTLKD----------LEEAFPECTYL   676
Baya_226_0010        PLDCFGDVVAKLPFSIPTEFNAGCLIIYLLMNPQSTFAE----------LSSVFPELYSL   679
LmjF.30.3430         PWDQFNNVVQRLPFNFPAEFSAGFLMTYVLCNPQCTMEE----------LCTTFPEMYAL   677
EMOLV88_300040300    PWDQFKNVVQLLPFNFPAEFSAGFLMTYVLCNPQCTMEE----------LCAVFPEMYAL   676

PCON_0044260         GKELEKVMAFTVHSLYLVKAYVRGHTAVVSEDLREDSYV----IDRALSLLSIKWEAHFD   1258
TCSYLVIO_007992      AVHLRMVMTFTLQALYLINAYKVSDKGTVVPQHLLEDNT----VECTIQLLWEKWYDHID   1197
TvY486_1001495       AKHLHIVMSFTLQALYLVNAYALTDREDIAPKDLLNGST----VAETIQMLWGKWRDHID   1348
Tb927.10.1490        ADHLHLVMSFTLQALYLINAYMLNDKETIVAKDQLTGTI----VEDTIEMMWQKWRDHID   1199
Baya_167_0010        PQQLRNVMEFTFQALYLLNAYRYYDHEPIRAQELTETSV----VEDTLGLMYEKWRLHFT   1259
LmjF.21.0820         QTQLRRVMEFTFHALYLLNVYILRDPSIVSSPELVETDV----VANALNLLHEKWVLHLD   1359
EMOLV88_210013000    QTQLRRVMEFTFHALYLLNVYILRDPSIVSSADLVETDV----VANALNLLHEKWVLHLD   1369
TCSYLVIO_003374      RHDLATLFYFWELAVCVLNSIACKDVP-LDVGCLHKANERMKELQRNLDINTGVRETYY-   734
TvY486_0604140       RHDLVTLFYFWDLAVQVLQRIGPKESSVSS-QHLSRANERMRRARNSLNIHAGMQETYS-   734
Tb927.6.4770         RHDLATLFYFWDLAVQVLQRIETKENFCVDQHCLSSANERMKRAQKNLNILTGVRETYY-   735
Baya_226_0010        EDDLRTLFWFWTMAFEAIVAIAQEEPDMVNLQGLLD-GHMHIS-DACSRVFPDFWAAMFA   737
LmjF.30.3430         DTDLQTLFWFFTMGFEAIRVINYEEPLSYDMVQVVNAARRLVG-QACERLCPDVFVAHFP   736
EMOLV88_300040300    DTDLQTLFWFFTMGFEAIRVINYEEPLSYDMVQVVNAARQLVG-QACERLCPEVYVENFP   735

PCON_0044260         SPFPEDTICGDLPYD--------------------DSEGAFSGNAM--------------   1284
TCSYLVIO_007992      EDPPQDIHGL-YGDS-----------------MIGMERGPAGG-----------------   1222
TvY486_1001495       DHPPQDIHNL-YTAG-----------------GCSS------------------------   1366
Tb927.10.1490        DNPPGDIHNL-YPPR-----------------HQEPIPH---------------------   1220
Baya_167_0010        DKPPSDVHQL-WEPE-----------------RSLES-----------------------   1278
LmjF.21.0820         GPAPEDVHGY-FQFE---------------------------------------------   1373
EMOLV88_210013000    GPAPTDVHGY-FEFE---------------------------------------------   1383
Baya_226_0010        PLQNTLNEAMMLNESL-LAAAQQQIPMQMPMMPPNHMSAGMGGI----------------   780
LmjF.30.3430         TADILFVAQ---NQGMDMGPMDPQMDVPM---PMNNMHGQMP-NMNMNMNMHPGMNSNMM   789
EMOLV88_300040300    TSDIIFVAQ---NQSMDMAPIDPQMDVPM---PMNNMHGQIPGNMNMNMNMHPSMNSNMM   789

Baya_226_0010        HY------ 782
LmjF.30.3430         QYQPPRMY 797
EMOLV88_300040300    QYQQPRMF 797
```
